# (p)ppGpp inhibits 70S ribosome formation in *Staphylococcus aureus* by impeding GTPase-ribosome interactions

**DOI:** 10.1101/2021.01.19.427108

**Authors:** Daniel J. Bennison, Jose A. Nakamoto, Timothy D. Craggs, Pohl Milón, John B. Rafferty, Rebecca M. Corrigan

## Abstract

During nutrient limitation, bacteria produce the alarmones (p)ppGpp as effectors of the stress signalling network termed the stringent response. Screening for (p)ppGpp-binding targets within *Staphylococcus aureus* identified four ribosome-associated GTPases (RA-GTPases), RsgA, RbgA, Era and HflX, each of which are cofactors in ribosome assembly, where they cycle between the ON (GTP-bound) and OFF (GDP-bound) states. Entry into the OFF-state from the ON-state occurs upon hydrolysis of GTP, with GTPase activity increasing substantially upon ribosome association. When bound to (p)ppGpp, GTPase activity is inhibited, reducing 70S ribosome assembly. Here, we sought to determine how (p)ppGpp impacts RA-GTPase-ribosome interactions by examining the affinity and kinetics of binding between RA-GTPases and ribosomes in various nucleotide-bound states. We show that RA-GTPases preferentially bind to 5′-diphosphate-containing nucleotides GDP and ppGpp over GTP, which is likely exploited as a regulatory mechanism within the cell. Binding to (p)ppGpp reduces stable association of RA-GTPases to ribosomal subunits compared to the GTP-bound state both *in vitro* and within bacterial cells by inducing the OFF-state conformation. We propose that in this conformation, the G2/switch I loop adopts a conformation incompatible with ribosome association. Altogether, we highlight (p)ppGpp-mediated inhibition of RA-GTPases as a major mechanism of stringent response-mediated growth control.

## Introduction

The prokaryotic 70S ribosome is an essential and complex macromolecular assembly responsible for the translation of messenger RNA (mRNA) into functional proteins. It comprises a large 50S and a small 30S subunit, which consist of 33 ribosomal proteins (r-proteins: L1-L36) associated with two ribosomal RNAs (rRNA), and 21 r-proteins (S1-S21) with one rRNA, respectively. Due to the energetic cost of ribosome synthesis and the intricacy of assembly, cofactors play a vital role in ensuring the correct conformation of the complete 70S (1). One class of assembly cofactors are the ribosome-associated GTPases (RA-GTPases), a subset of P-loop GTPases within the Translation Factor Associated (TRAFAC) family, of which the proteins RsgA, RbgA, Era and HflX are members. RA-GTPases have a highly conserved G-domain housing the catalytic G1-G5 motifs (Supplementary Figure S1), flanked by one or more highly variable accessory domains that convey targeting and additional functionality to the enzymes (Figure 1A) (2-6). The high degree of sequence identity (Supplementary Figure S1A) and structural conservation (Supplementary Figure S1B-E) between functional motifs within the nucleotide-binding pocket suggests a consistent mechanism of binding among these P-loop RA-GTPases.

**Figure 1.**
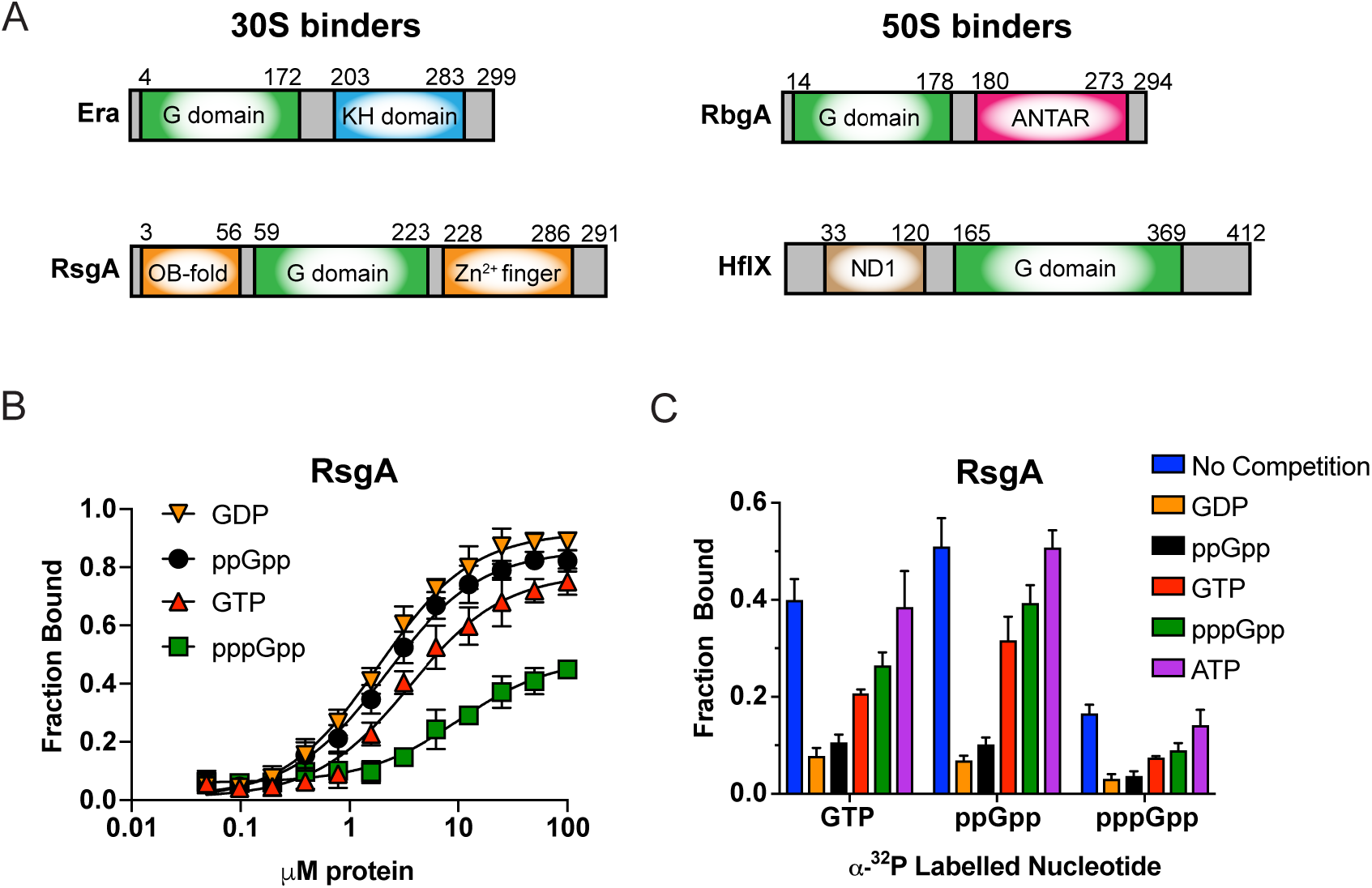
Nucleotide binding characteristics of RA-GTPases. A) Schematic representation of the domain structure of Era, RsgA, RbgA and HflX from *S. aureus*. The conserved GTPase domain (G domain) is coloured in green and accessory domains are shown. B) Determination of binding affinities and *K*_d_ values for ^32^P-labelled GTP, GDP, ppGpp and pppGpp with purified recombinant RsgA using DRaCALA as previously described (36). Each point is the mean average of at least 3 technical replicates and error bars indicate standard deviation. C) DRaCALA binding assay of recombinant RsgA binding to ^32^P-labelled GTP, ppGpp and pppGpp in the presence of an excess of cold competitor (GTP, GDP, ppGpp, pppGpp or ATP). All experiments were carried out in triplicate, with error bars representing standard deviation.

Due to the variation in accessory domains, each RA-GTPase associates with a distinct area of the ribosome to coordinate a maturation event. Cycling between the GTP-bound ON and GDP-bound OFF states enables these proteins to act as molecular checkpoints of ribosome assembly by monitoring the maturation state of individual subunits (7). Although it is unclear what the precise roles of RA-GTPases are in ribosomal maturation, they have been suggested to sterically prevent the premature association of other r-proteins (8). Unknown maturation events then act as activators of GTPase activity, enabling entry into the GDP-bound OFF state and subsequent dissociation from the ribosome (7). In addition to regulating the recruitment of r-proteins, RA-GTPases have been postulated to recruit RNA processing enzymes directly. For instance, the RA-GTPase Era can interact with several proteins involved in 16S rRNA maturation, including YbeY, an endonuclease involved in 16S processing in *Escherichia coli* (9), and CshA, a DEAD-box RNA helicase (10), pointing to a role for this group of enzymes as hub proteins that facilitate maturation events.

During periods of starvation, bacteria produce the alarmones guanosine penta- and tetraphosphate (collectively referred to as (p)ppGpp), which function as the mediators of a stress signalling system termed the stringent response (11). Amidst this response, the concentration of (p)ppGpp within the cell can reach between 1-2 mM with a concurrent drop in GTP levels (12,13). This results in a plethora of downstream effects, including alterations to transcription, translation and DNA replication, as well as regulating late-stage growth phases such as sporulation or biofilm formation (14-16). Our previous work identified the four RA-GTPases (RsgA, RbgA, Era and HflX: Figure 1A) in the pathogenic bacterium *Staphylococcus aureus* as enzymes that can bind to and are inhibited by (p)ppGpp, resulting in a negative impact on 70S ribosome assembly and growth (17). RsgA is a nonessential, highly conserved late-stage 30S assembly cofactor (17,18), that has been implicated in the docking of helix 44 (h44) of the 16S rRNA into the correct conformation and therefore correct maturation of the decoding centre prior to subunit joining (4,19). Era is a highly conserved protein, known to interact with the 3’ end of the pre-16S rRNA (3) where it monitors the ribonuclease processing state of this region. Furthermore, since Era docking occurs at the same site as r-protein S1 adjacent to the anti-Shine-Dalgarno sequence, it can also sterically occlude initiation factor 3 (IF3) binding and hence prevent formation of the 30S pre-initiation complex (20). RbgA is a late-stage 50S binding protein, implicated in RNA binding and remodelling (6,21). Finally, HflX is a 30S, 50S and 70S binding protein that has been implicated in the splitting and subsequent repair of heat-stalled 70S ribosomes (22). HflX also contributes directly to 70S levels through GTPase-dependent splitting of the 100S hibernation complex to enable rapid recovery of active 70S ribosomes when cellular energy levels rise, a process that is inhibited when bound to (p)ppGpp (23).

The binding of pppGpp to RbgA has previously been suggested to enhance the affinity of this protein for the mature 50S subunit when compared to the GTP-bound form (24). More recently the crystal structure of *S. aureus* RbgA in complex with both ppGpp and pppGpp was solved, revealing a competitive mode of inhibition at the catalytic centre (6). These findings have led to a proposed model where RbgA-(p)ppGpp likely sequesters 50S ribosomal subunits to prevent formation of active 70S ribosomes (6). Here, we further characterise the four RA-GTPases RsgA, RbgA, Era and HflX, to investigate the relationship between RA-GTPases and stringent response-mediated control of ribosome assembly in *S. aureus*. We find that the 5’ diphosphate nucleotides GDP and ppGpp can bind to these enzymes with higher affinity than the 5’ triphosphate-containing GTP or pppGpp, suggesting that occupancy of the binding site is strongly dependent on a cellular excess of GTP over GDP, which occurs in exponential and non-stressed cells (25). Contrary to previous models, we demonstrate that interactions with (p)ppGpp prevent the association of RA-GTPases to the ribosome, both *in vitro* and in *S. aureus*. To examine mechanistically how (p)ppGpp can prevent RA-GTPase-ribosome interactions we use X-ray crystallography, revealing that (p)ppGpp binding causes the RA-GTPases to adopt a conformation similar to the inactive GDP-bound OFF state, with the switch I/G2 loop required for GTP hydrolysis extended away from the catalytic site where it could sterically hinder interactions with the ribosome. Altogether, we propose a mechanism behind (p)ppGpp-controlled inhibition of ribosome assembly and increase our understanding of stringent response-mediated translational control by means of RA-GTPase inhibition.

## MATERIALS AND METHODS

### Bacterial strains and culture conditions

*E. coli* strains were grown in Luria Bertani broth (LB) and *S. aureus* strains in tryptic soy broth (TSB) at 37°C or 30°C with aeration. Strains and primers are listed in Supplementary Tables S1 and S2. Antibiotics were used when appropriate at the following concentrations: kanamycin 30 µg/ml, carbenicillin 50 µg/ml, spectinomycin 250 µg/ml and tetracycline 2 µg/ml. pET28b-*era* G2 was created using overlap primer PCR with plasmid pET28b-*era* as a template and primers as listed in Supplementary Table S2. pCN55iTET-*era*-His was constructed by amplifying *era* using primers RMC157/RMC536 from LAC* genomic DNA and cloning into the KpnI/SacI sites of pCN55iTET. All plasmids were initially transformed into *E. coli* strain XL1-Blue and sequences of all inserts were verified by fluorescence automated sequencing by GATC. For protein expression and purification, all pET28b derived plasmids were transformed into *E. coli* strain BL21 (DE3). All *S. aureus* plasmids were first electroporated into RN4220 Δ*spa*, before isolation and electroporation into LAC* Δ*era*.

### GTPase assays

GTPase activity assays were performed as previously described (10). Briefly, the ability of proteins to hydrolyse GTP was determined by incubating 100 nM recombinant protein with 100 nM *S. aureus* 70S ribosomes, 1 μM GTP and 2.78 nM α-^32^P-GTP in 40 mM Tris pH 7.5, 100 mM NaCl (100 mM KCl for RbgA), 10 mM MgCl_2_ at 37°C for the indicated times. All reactions were also set up in the absence of enzymes to monitor spontaneous GTP hydrolysis. Reactions were heat inactivated at 95°C for 5 mins to precipitate proteins and release bound nucleotide. Precipitated proteins were pelleted by centrifugation at 17,000 x g for 10 min. Reaction products were visualized by thin layer chromatography (TLC) in PEI cellulose TLC plates (Macherey-Nagel) and separated using 0.75 M KH_2_PO_4_, pH 3.6 buffer. The radioactive spots were exposed to a BAS-MS Imaging Plate (Fujifilm), visualised using an LA 7000 Typhoon PhosphorImager (GE Healthcare), and images quantified using ImageQuant (GE Healthcare).

### Synthesis of ^32^P-(p)ppGpp, differential radial capillary action of ligand assays (DRaCALA)

The synthesis of (p)ppGpp and DRaCALA binding and competition assays were performed as described previously (17).

### Protein purifications

Proteins were purified from 1-2 L *E. coli* BL21 DE3 cultures. Cultures were grown at 37°C to an OD_600_ of 0.5-0.7, expression was induced with 1 mM isopropyl β-D-1-thiogalactopyranoside (IPTG) and incubated for 3 h at 30°C. Cell pellets were resuspended in 5 ml Buffer A (50 mM Tris pH 7.5, 150 mM NaCl, 5% glycerol, 10 mM imidazole) and lysed by sonication upon addition of 20 μg/ml lysozyme and 30 μg/ml RNase A. Protein purifications were performed by nickel affinity chromatography. The filtered cell lysate was loaded onto a 1 ml HisTrap HP Ni^2+^ column (GE Healthcare) before elution using a gradient of Buffer B (50 mM Tris pH 7.5, 200 mM NaCl, 5% glycerol, 500 mM imidazole). Protein containing fractions were dialysed in 50 mM Tris-HCl pH 7.5, 200 mM NaCl, 5% glycerol before concentration using a 10 kDa centrifugal filter (Thermo Scientific) and storage at-80°C. Protein for use in crystallography was dialysed into 25 mM Tris-HCl pH 7.5, 200 mM NaCl and used immediately. Protein concentrations were determined by absorbance at 280 nm using appropriate extinction coefficients. A_260/280_ ratios were monitored to ensure preparations had low RNA/nucleotide contamination (<5%), indicated by a ratio below 0.8. The extinction coefficients at 280 nm for each protein and their mutant variants were calculated from the primary structure: Era: 25900 M^-1^ cm^-1^, RsgA: 23505 M^-1^ cm^-1^, RbgA: 40910 M^-1^ cm^-1^, HflX: 24870 M^-1^ cm^-1^. Typically, protein purity was above 95% as assayed by 12% SDS-PAGE and Coomassie-blue staining.

### 30S, 50S and 70S ribosome purification

70S ribosomes were purified as described (17), with the following exceptions: following purification of mature 70S ribosomes, the ribosome pellet was resuspended in dissociation buffer (20 mM Tris pH 7.5, 120 mM NH_4_Cl, 1.5 mM MgCl_2_ and 2 mM β-mercaptoethanol), and quantified using the absorbance at 260 nm as described (26). 50 A260 units of 70S ribosomes were applied to a 10-40% continuous sucrose gradient made up in dissociation buffer and separated at 111,000 x g for 16 hours. Gradients were fractionated by upwards displacement of 250 µl aliquots, which were analysed for RNA content at an absorbance of 260 nm. Fractions containing 30S and 50S ribosomal subunits were pooled separately, and purification was continued as described (26).

### *In vitro* ribosome association assays

500 nM recombinant 6xHis-tagged RA-GTPase was incubated at room temperature for 5 mins with 200 nM *S. aureus* 70S ribosomes in dissociation buffer (20 mM Tris pH 7.5, 120 mM NH_4_Cl, 1.5 mM MgCl_2_ and 2 mM β-mercaptoethanol) in the apo form and in the presence of 40 µM GTP, GMPPNP, GDP, ppGpp or pppGpp. The resultant reaction (150 μl) was layered onto a 10-40% continuous sucrose density gradient in dissociation buffer. Subsequently, gradients were centrifuged for 16 h at 111,000 x g in order to separate the 30S and 50S subunits. Gradients were fractionated by upwards displacement of 250 µl aliquots, which were analysed for RNA content at an absorbance of 260 nm. Fractions containing 30S and 50S ribosomal subunits were pooled separately and the protein content was precipitated by the addition of 10% v/v trichloroacetic acid (TCA) and incubation for 3 h at 4°C. Samples were centrifuged at 17,000 x g for 5 mins and washed twice with ice-cold acetone prior to drying of the pellets at 37°C for 10 mins. Pellets were resuspended in 2x SDS-PAGE sample buffer (62.5 mM Tris-HCl pH 6.8, 2% SDS, 10% glycerol, 0.01% bromophenol blue, 10% v/v β-mercaptoethanol), proteins were separated using a 10% SDS-PAGE gel and transferred onto a PVDF Immobilon-P membrane (Merck Millipore). The membrane was blocked with 5% w/v milk in TBST (50 mM Tris-HCl pH 7.6, 150 mM NaCl, 0.1% Tween 20), probed using 1:500 monoclonal anti-His HRP-conjugated antibodies (Sigma) and imaged using a ChemiDoc MP (Bio-Rad). Band densitometry was performed using ImageJ.

### Growth and *in vivo* ribosome association assays

*S. aureus* strains were grown overnight in TSB containing the appropriate antibiotics. Overnight cultures were diluted to a starting OD_600_ of 0.05 in the presence of 100 ng/ml Atet and appropriate antibiotics and grown at 37°C with aeration, with OD_600_ values determined at 2 h intervals. For ribosome association assays, a culture of LAC* *Δera* pCN55iTET-*era*-his was split at an OD_600_ of 0.6 and fractions were either left uninduced or were induced with either 0.05 or 60 µg/ml mupirocin at 37°C for 30 mins. After growth, all cultures were incubated with 100 μg/ml chloramphenicol at 37°C for 3 mins, then cooled to 4°C. Cells were centrifuged at 4,000 x g for 10 mins and pellets resuspended to an OD_600_ of 35 in dissociation buffer (20 mM Tris pH 7.5, 120 mM NH_4_Cl, 1.5 mM MgCl_2_ and 2 mM β-mercaptoethanol). Cells were lysed through the addition of 0.5 µg/ml lysostaphin and 75 ng/ml DNase for 60 mins at 37°C. Lysates were centrifuged at 17,000 x g for 10 min to remove cell debris and 250 µl of the lysate was layered onto a 10-40% continuous sucrose gradient in dissociation buffer. Subunit separation was continued as per the *in vitro* method and associated C-terminally histidine-tagged Era (Era-His) was quantified via western blotting and band densitometry (ImageJ). Crude lysates were loaded alongside pulled-down protein to verify Era-His expression level. Staining of the blotting membrane with Ponceau S in 5% acetic acid was used to ensure consistent lysate loading prior to membrane blocking. Membranes were incubated with staining solution for up to 5 minutes and washed with distilled water until the background was clear. Following imaging, the Ponceau S was removed by repeated wash steps using PBS.

### Ribosome profiles from *S. aureus* cell extracts

Crude isolations of ribosomes from *S. aureus* cell extracts were achieved as described by Loh *et al*. with some modifications (27). Briefly, 100 ml cultures of the different *S. aureus* strains were grown to an OD_600_ of 0.4 in TSB medium with 100 ng/ml anhydrotetracycline (Atet). 100 μg/ml chloramphenicol was added to each culture and incubated for 3 min before being cooled to 4°C to enhance the pool of 70S ribosomes. Pelleted cells were suspended in association buffer (20 mM Tris-HCl pH 7.5, 8 mM MgCl_2_, 30 mM NH_4_Cl and 2 mM β-mercaptoethanol) and normalized to an OD_600_ of 15. Cells were lysed by the addition of 0.2 μg/ml lysostaphin and 75 ng/ml DNase and incubated for 60 min at 37°C. Cell debris was removed by centrifugation at 17,000 x g for 10 min. Clarified lysates (250 μl) were layered onto 10-50% discontinuous sucrose density gradients made in association buffer. Gradients were centrifuged for 7 h at 192,100 x g. Gradients were fractionated by upwards displacement of 250 μl aliquots, which were analysed for RNA content by absorbance at 260 nm.

### Crystallisation of RsgA

The purified recombinant protein consisted of 311 residues, comprising 291 residues of *S. aureus* RsgA with an N-terminal 20 residue tag MGSSHHHHHHSSGLVPRGSH. It was simultaneously buffer exchanged into 25 mM Tris-HCl pH 7.5, 200 mM NaCl buffer and concentrated to 30 mg/ml for crystallization screening using the sitting drop vapour diffusion method. Each droplet contained 200 nl protein solution and 200 nl crystallisation reagent from an adjacent well of 50 µl volume. Figures were prepared in Pymol (The PyMOL Molecular Graphics System, Version 2.0 Schrödinger, LLC), with the exception of electron density maps which were generated using COOT (28,29).

#### RsgA-ppGpp

The concentrated RsgA solution was supplemented with 2 mM MgCl_2_ and 2 mM ppGpp. Successful crystallisation was observed when this sample was mixed 1:1 with well solution containing 0.2 M sodium citrate tribasic dihydrate, 0.1 M Bis-Tris propane pH 6.5 and 20% (w/v) PEG 3350, and incubated at 17°C. Rod shaped crystal clusters appeared after a few days. Crystals were transferred to a cryoprotectant solution consisting of mother liquor with 15% ethylene glycol added and flash cooled in liquid N_2_. X-ray diffraction data were collected from a single crystal on beamline i04 at the Diamond Light Source national synchrotron facility at a wavelength of 0.97949 Å. The ppGpp-bound crystals diffracted to a resolution of 1.94 Å (PDB: 6ZHL). Initial processing was completed using the Xia2 pipeline (30). The crystals belonged to the space group P2_1_2_1_2_1_ (Supplementary Table S3). The structure of RsgA-ppGpp was solved via molecular replacement, using the previously published *Bacillus subtilis* homologue YloQ (PDB: 1T9H) as a model. The structure contained one RsgA monomer in the asymmetric unit. Molecular replacement was carried out using Phaser from within the CCP4 suite (31,32). The structure was refined via rounds of manual model building and refinement using COOT (29) and REFMAC5 (33). The final model was validated using MOLPROBITY (34). Residues 181-200 were lacking electron density and as such were omitted from the final model.

#### Apo RsgA

Crystallisation of apo RsgA was achieved when the concentrated protein sample was mixed 1:1 with well solution containing 0.15 M ammonium sulphate, 0.1 M MES pH 6.0 and 15% (w/v) PEG 4000 and incubated at 17°C. A single rod shaped crystal formed after a few weeks and diffracted to 2.01 Å resolution (PDB: 6ZJO). Initial processing was completed using the Xia2 pipeline and the crystal belonged to the space group P12_1_1 (Supplementary Table S3). The structure was solved via molecular replacement as above using the available RsgA-ppGpp structure as a model with ligands removed and contained two RsgA monomers in the asymmetric unit. Iterative rounds of modelling, refinement and validation were carried out as above. Residues 180-200 (Chain A) and 179-200 (Chain B) were lacking electron density and as such omitted from the model.

### Fluorescent labelling of proteins

200 µM recombinant protein was incubated with 5 mM dithiothreitol (DTT) for 1 h at room temperature. DTT was removed via two consecutive passes through a PD-10 Sephadex G-25 M buffer exchange column (GE Healthcare) as per the manufacturer’s instructions into labelling buffer (50 mM HEPES pH 7.1, 200 mM KCl, 5% glycerol, 120 µM TCEP). Flow-through was analysed for protein content at 280 nm. 50 µM of reduced protein was incubated with 100 µM ATTO 488-maleimide (ATTO-TEC) overnight at 4°C, shielded from light and subject to gentle shaking. The reaction was stopped by addition of 6 mM β-mercaptoethanol and mixtures were applied to a 1 ml HisTrap HP Ni^2+^ column (GE Healthcare) before elution using a gradient of Buffer B (50 mM Tris pH 7.5, 200 mM NaCl, 5% glycerol, 500 mM imidazole) and subsequent dialysis to remove imidazole. Labelling efficiency was calculated in accordance with the fluorescent dye manufacturer’s guidelines.

### Stopped-flow fluorescence kinetics measuring ribosome association

For initial controls, 0.2 µM Atto-488-labelled proteins were rapidly mixed with 0.2 µM *E. coli* 50S ribosomal subunits (purified as described elsewhere (35)) in TAKM7 buffer (25 mM Tris-HCl pH 7.4, 70 mM ammonium acetate, 30 mM KCl and 7 mM MgCl_2_) using an SX20 stopped-flow apparatus (Applied Photophysics) in the presence or absence of GTP, ppGpp and pppGpp. Equal volumes (60 µl) of each reactant were rapidly mixed at 25°C. Atto-488 was excited using a 470 nm LED and fluorescence was detected through a 515 nm long-pass filter. Reactions were monitored for 10 seconds using a logarithmic sampling method, with 1000 total datapoints per reaction. Each condition was subject to at least 5 technical repeats, with curves representing the mean average fluorescence of the technical repeats.

For titrations, 0.075 µM RbgA or 0.05 µM HflX labelled proteins were mixed with a 200-fold excess of GTP, ppGpp or pppGpp (15 µM for RbgA and 10 µM for HflX) in TAKM7 buffer just prior to use. *E. coli* ribosomal 50S subunits were used in excess relative to the labelled protein in the presence of nucleotides, up to 0.8 µM. Samples were then loaded separately into an SX20 stopped flow apparatus. Equal volumes (60 µl) of each reactant were rapidly mixed at 25°C and fluorescence emission was monitored as described above. The resultant fluorescence timecourses were fitted using the double exponential function 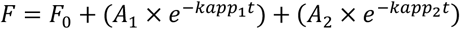 with fluorescence signal at time *t* (*F*), initial fluorescence signal (*F*_0_), the amplitude of signal change of the first exponential (*A*_1_), the apparent rate of the first exponential (*k*_app1_), the amplitude of signal change of the second exponential (*A*_2_), the apparent rate of the second exponential (*k*_app2_) and time (*t*). Each timecourse was fitted individually, with curves shown representing the mean average of at least 5 technical replicates. If necessary, a linear term was included. Data was normalised to the mean of the first 10 fluorescence measurements. The microscopic constants *k*_1_, *k*_-1_, *k*_2_ and *k*_-2_ were calculated by plotting both the sum and product of the apparent rates *k*_app1_ and *k*_app2_ for each titration and analysing the resulting linear relationship using linear regression. Briefly, taking *A* as the linear regression of the sum of *k*_app1_ and *k*_app2_, and *B* as the linear regression of the product of *k*_app1_ and *k*_app2_, kinetic parameters were determined as follows: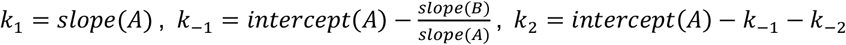, and 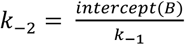. Dissociation constants (*K*_d_) were calculated using the following equation: 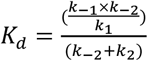.

### Statistics

Statistical analyses were performed using Graphpad Prism 8.0 software. Statistical differences between samples were assessed using one-way analysis of variance (ANOVA), followed by Tukey’s multiple comparisons test.

## RESULTS

### RA-GTPases preferentially bind 5’ diphosphate-containing nucleotides GDP and ppGpp

The RA-GTPases RsgA, Era, RbgA and HflX can bind to the guanosine nucleotides GTP, GDP, ppGpp and pppGpp. Our previous work observed higher binding affinities for ppGpp over GTP, pointing towards a difference in binding between 5’ di- or triphosphate nucleotides (17). To examine the nucleotide binding affinities of these RA-GTPases for GDP in comparison to ppGpp, pppGpp and GTP, we used a differential radial capillary action of ligand assay (DRaCALA) (Figure 1B, Supplementary Figure S2A-C, (36)). In each case, the affinities of 5’ diphosphate-containing GDP and ppGpp were similar in the low µM range and were 2-6-fold higher than the affinities of either GTP or pppGpp (Supplementary Table S4). This supports a previous observation that ppGpp is a more potent inhibitor of GTPase activity than pppGpp (17).

Structural data places (p)ppGpp within the GTP-binding site of the RA-GTPase RbgA (6). Based on our measured affinities (Supplementary Table S4), we speculate that both GDP and ppGpp will out-compete other nucleotides for occupancy of the binding site. To examine this, competition assays were performed in which the binding of a radiolabelled nucleotide was challenged with an excess of unlabelled nucleotides (Figure 1C, Supplementary Figure S2D-F). In each case, addition of cold unlabelled nucleotide reduced occupancy of the labelled nucleotide, with the exception of the ATP control. This is likely due to the much lower affinity of RA-GTPases for adenosine bases conveyed by a contact from the conserved aspartate residue of the G4 motif to the 2-amino group of the guanosine base (37). A hierarchy of binding could be established depending on the level of competition provided by each unlabelled nucleotide, with GDP and ppGpp competing more effectively (Figure 1C, Supplementary Figure S2D-F). This suggests that GTP occupancy, and hence activity, of these RA-GTPases is strongly dependent on the cellular excess of GTP over GDP and ppGpp, which occurs during exponential growth when ribosomal biogenesis is at its peak (25). This ratio changes during stationary phase and upon induction of the stringent response, when cellular GTP levels decrease with a concurrent rise in (p)ppGpp (12,38), shifting binding to favour a ppGpp-bound state. The greater affinity of these RA-GTPases to diphosphate-containing nucleotides would hence aid a rapid transition between the GTP-bound and ppGpp-bound states under conditions of stress.

### Interactions with (p)ppGpp reduce the affinity of RA-GTPases for the ribosome

It is well characterised that rRNA transcription decreases during the stringent response (39). In addition, the GTPase activity of ribosome assembly cofactors is inhibited by (p)ppGpp, both of which contribute to a reduction in mature ribosomes within the cell (17). To examine mechanistically how (p)ppGpp-GTPase interactions affect the ability of RA-GTPases to associate with ribosomal subunits, we examined the association of each GTPase to either the 30S or 50S ribosomal subunit in the presence of GDP, GTP, ppGpp, pppGpp, as well as GMPPNP, a non-hydrolysable analogue of GTP. His-tagged GTPases were preincubated with highly pure, salt washed 70S *S. aureus* ribosomes in a low magnesium buffer to encourage ribosomal subunit dissociation, and the amount of each GTPase associated with each of the subunits was quantified by western immunoblot using anti-His antibodies after sucrose gradient separation (Figure 2). In all cases, we observed a marked decrease in association of each GTPase to the 30S or 50S subunits in the presence of GDP, ppGpp and pppGpp compared to the GMPPNP-bound state (Figure 2A-D). For Era and HflX, there was a similar level of subunit association when in the apo, GTP or GMPPNP-bound states, compared to a 2-fold reduction in ribosome binding when incubated with GDP, ppGpp or pppGpp (Figure 2C, 2D), suggesting that these GTPases can associate to the ribosome in the unbound state. The ability of Era to bind the 30S in the absence of nucleotides has been reported previously, where it has been suggested that the apo form can bind to a secondary site (3,40). The patterns exhibited by RsgA and RbgA were slightly different, with strong binding in the GMPPNP-bound state, whereas 3-6 fold weaker binding was observed in the apo, GTP, GDP, ppGpp and pppGpp-bound states (Figure 2A, 2B). It is worth noting that previous studies have suggested that the association of RbgA with the 50S subunit is enhanced in the presence of pppGpp (24), a finding that is not replicated here. The apparent effect of ppGpp and pppGpp on ribosome association was comparable, which is not reflective of the differences in affinity (Figure 1B, Supplementary Figure S2A-C), although under the conditions tested here the excess of nucleotide would maintain an equilibrium favouring the nucleotide-bound state. Furthermore, the four RA-GTPases were found to be unable to hydrolyse pppGpp, and as such conversion of pppGpp to ppGpp was not responsible for the similar degree of inhibition of association. We postulate that the low level of binding observed when preincubated with GTP could be due to GTP hydrolysis during the 16 hour centrifugation step, likely causing the GTPases to enter the GDP-bound state and dissociate. This, in turn, may be enhanced by the higher affinity of GDP for these GTPases as compared to GTP (Supplementary Table S4). From these data, we show that GTP binding favours association of RsgA, RbgA, Era and HflX to ribosomal subunits, and that this interaction is inhibited when in the GDP-, ppGpp- or pppGpp-bound states.

**Figure 2.**
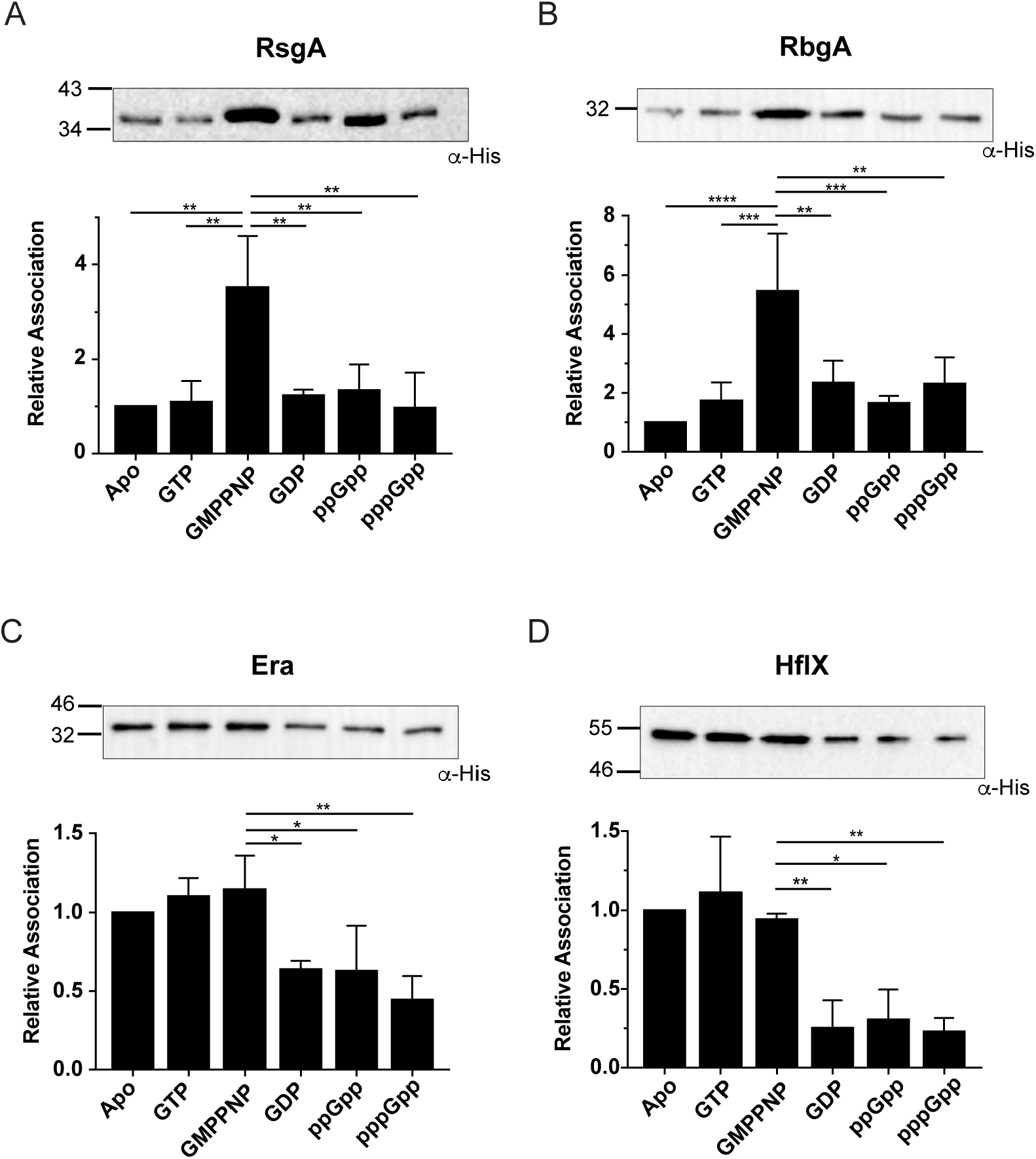
(p)ppGpp and GDP binding reduces RA-GTPase association to the ribosome. A-D) Top: purified 70S ribosomes were preincubated with His-tagged A) RsgA, B) RbgA, C) Era and D) HflX in the absence or presence of GTP, GMPPNP, GDP, ppGpp or pppGpp. Following subunit separation and precipitation, bound proteins were detected using HRP-conjugated α-His antibodies. Experiments were carried out in triplicate or quadruplicate and one representative image of each are shown. Bottom: the signal intensities relative to the apo-state of all repeats are plotted with error bars representing standard deviation. Statistical analysis was performed using a one-way ANOVA followed by Tukey’s multiple comparison test. (* *P* < 0.05, ** *P* < 0.01, *** *P* < 0.001, **** *P* < 0.0001).

### Binding kinetics of RA-GTPase-ribosome interactions

To gain further insight into the binding mechanism and how (p)ppGpp reduces the association of RA-GTPases to the ribosomal subunits, we used a stopped-flow technique with fluorescent derivatives of the RA-GTPases (Figure 3A). Structural predictions of all four R A-GTPases were built by homology modelling using available structures to assess the availability of suitable residues for fluorescence labelling (Supplementary Figure S3A, S3B)(41). Both RbgA and HflX were amenable to covalent linkage to the fluorophore Atto-488 using maleimide chemistry with exposed cysteine residues. RbgA contains one wild-type cysteine residue (C277) that is surface exposed in the *B. subtilis* crystal structure (PDB: 1PUJ) and is located towards the C-terminus of the protein (Supplementary Figure S3A). Based on the *E. coli* structure (PDB: 5ADY), HflX contains two cysteines (Supplementary Figure S3B). C330 is predicted to be surface exposed and therefore amenable to labelling, while C45 is buried and is expected to show low accessibility for fluorescent labelling. Era, on the other hand, lacks any cysteine residues, while RsgA contains three conserved cysteine residues that coordinate the Zn^2+^ ion within the Zn^2+^-finger domain (ZNF), and as such both were not suitable for labelling. Both Atto488-labelled RbgA and HflX retained wild-type levels of GTPase activity, which can still be inhibited by ppGpp (Supplementary Figure S3C, S3D).

**Figure 3.**
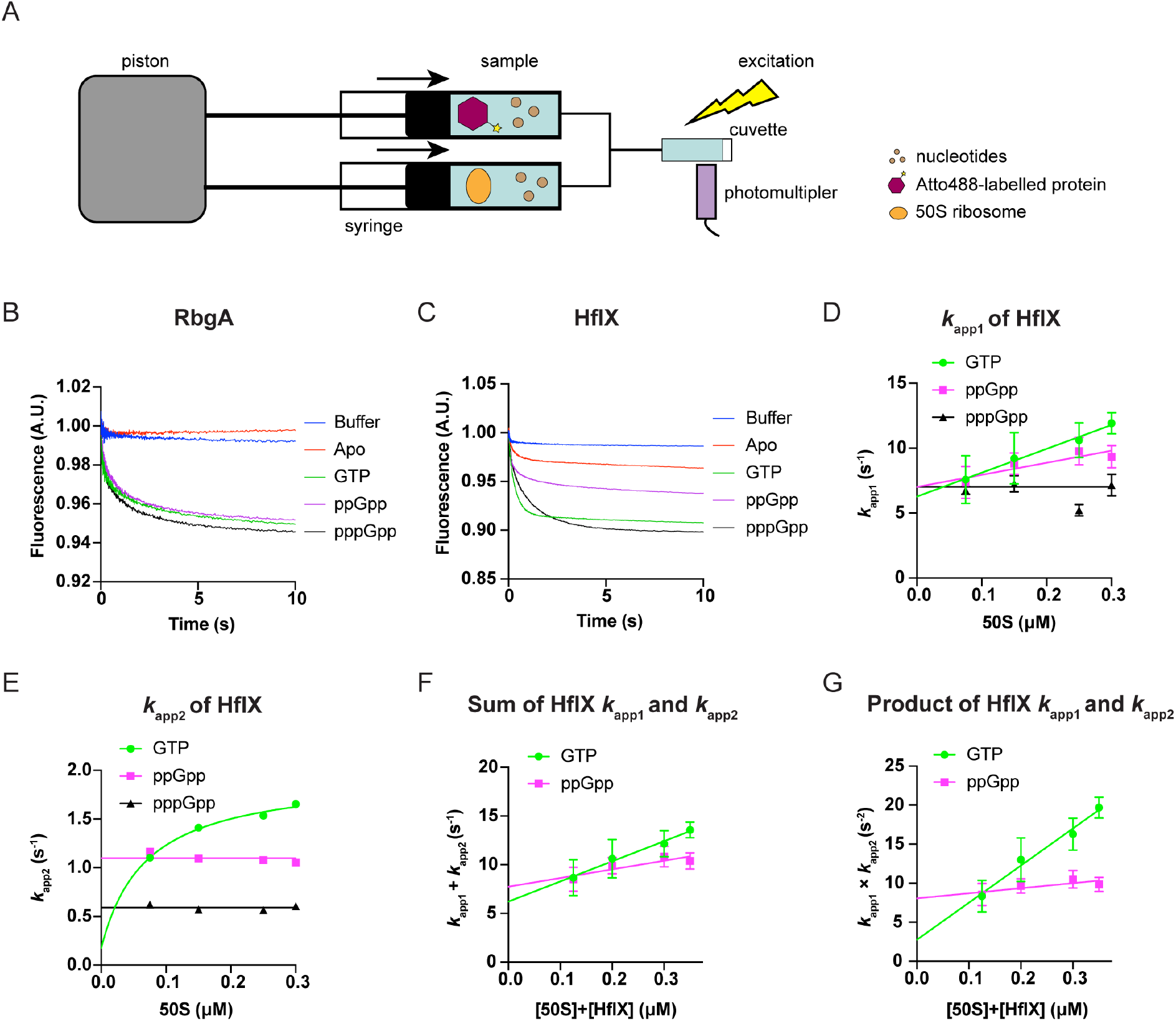
Stopped-flow kinetic parameters of RA-GTPase association to the ribosomal subunits. A) Schematic representation of the experimental setup for stopped-flow analysis. Nucleotides (brown circles), 50S subunits (orange oval) and Atto488-labelled RA-GTPases (purple hexagon) are indicated. Arrows indicate the direction of syringe movement. Atto-488 was excited using a 470 nm LED and fluorescence was detected through a 515 nm long-pass filter. B, C) Fluorescent change upon mixing 0.2 µM RbgA-Atto488 (B) or HflX-Atto488 (C) with 0.2 µM 50S ribosomal subunits in the presence of 100 µM GTP, ppGpp and pppGpp or in the Apo state using the stopped flow fluorescence apparatus. Fluorescently labelled protein was also mixed with buffer lacking 50S subunits as a mixing control. Fluorescence of the reaction was tracked using exponential sampling for 10 seconds and each curve is the mean average of at least 5 technical replicates. D) *k*_app1_ dependence on 50S concentration for HflX complexed with GTP (green), ppGpp (pink), pppGpp (black). E) as (D) for the *k*_app2_ dependence. F) and G) Sum and product analyses of apparent rates during HflX association to the 50S subunit. 0.05 µM HflX-Atto488 was mixed with increasing titrations of 50S ribosomal subunits over the fluorescently labelled protein, in the presence of 20 µM GTP or ppGpp. The resultant traces (Supplementary Figure S4) were analysed by nonlinear regression using two exponential terms. The sum (F) and product (G) of apparent rates (*k*_app1_ (D), *k*_app2_ (E)) were plotted as a function of the total concentration of the 50S subunits and HflX protein to determine the microscopic constants *k*_1_, *k*_-1_, *k*_2_, and *k*_-2_ (Supplementary Table S5) and the resulting dissociation constant (*K*_d_) (see Methods section). Error bars represent the standard deviation of the apparent rates of 4 or more individual traces (D, E) or the standard error of the two-step analysis (F, G).

Using the fluorescent variants of RbgA and HflX, we studied the binding mechanism of both to the 50S ribosomal subunit in the GTP-, ppGpp- and pppGpp-bound states. First, the fluorescence change of each labelled protein was measured upon the interaction with activated mature ribosomal subunits in the presence of different nucleotides (Figure 3B, 3C). RbgA showed no change in fluorescence while in the apo state, indicating a lack of interaction with the ribosome. On the other hand, all nucleotide-bound states showed a large decrease in fluorescence when mixed with the 50S subunit, consistent with some level of 50S association taking place when bound to GTP, ppGpp or pppGpp (Figure 3B). HflX, on the other hand, exhibited a fluorescence change upon mixing with the 50S subunit in the absence or presence of all tested nucleotides (GTP, ppGpp and pppGpp), which could be taken as a direct measure of ribosome association changing the chemical environment of the fluorophore (Figure 3C).

Next, we used a constant concentration of protein in the presence of 200-fold excess of each nucleotide and titrated it with increasing concentrations of ribosomal subunits (Supplementary Figure S4). Time traces appeared biphasic for both RA-GTPases independent of the nucleotide bound. Analysis of the fluorescent time traces with a double exponential equation yielded the apparent rates of association (*k*_app1_, *k*_app2_) (Figure 3D, 3E, Supplementary Figure S5A, S5B), in accordance with a binding mechanism composed of two sequential steps. Thus, the mechanism describing the interaction (Equation 1: 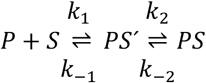) consists of an initial interaction and subsequent stabilisation of the factor on the ribosome, where *P* is the protein, *S* is the ribosomal subunit, *PS’* is the transient complex and *PS* is the stable complex.

For two-step reactions, the apparent rate under conditions tested, *k*_app1_, is expected to increase linearly with increasing ligand concentration. On the other hand, *k*_app2_ is expected to align to a hyperbolic relationship as ligand concentration increases. This was the case for HflX complexed with GTP (Figure 3D, 3E). Thus, productive binding of the RA-GTPase appears to occur through two steps. When HflX was incubated with ppGpp, the *k*_app1_ increased linearly (Figure 3D), while *k*_app2_ did not depend on ribosome concentration (Figure 3E), indicating that ppGpp hampers the accommodation step of the binding mechanism. On the other hand, if HflX was complexed with pppGpp, neither *k*_app_ depended on 50S concentration, indicating that the alarmone drastically affects the mechanism of HflX binding. In this case, the reaction appears rate-limited by an isomerization step of the RA-GTPase at 5 s^-1^ (Figure 3D). The linear increase in *k*_app1_ was 2-fold greater for GTP than for ppGpp or pppGpp (Figure 3D), suggesting a greater rate of the fast-phase reaction. The *k*_app2_ of the GTP-bound form showed a hyperbolic relationship tending to 2 s^-1^, while the linear relationship when bound to ppGpp was steady at 1.0 s^-1^ (Figure 3E). This suggests that the second, slow-phase reaction is taking place while HflX is bound to GTP but is reduced 4-fold when bound to ppGpp. Additionally, this suggests that one or more of the microscopic constants which contribute to the *k*_app2_ in the two-step association reaction remains incomplete while in the ppGpp-bound state.

Next, we used the sum and product of the *k*_app1_ and *k*_app2_ of each reaction (Figure 3F, 3G) to estimate the microscopic constants defining the reaction for the GTP- and ppGpp-bound HflX (Supplementary Table S5). ppGpp reduced the value of the initial binding constant *k*_1_ by 2-3 fold, while drastically affecting *k*_2_, indicating that the alarmone hampers proper accommodation of HflX on the subunit (Figure 3F, 3G, Supplementary Table S5). On the contrary, the dissociation rate constants *k*_-1_ and *k*_-2_ appeared less affected by ppGpp, remaining similar to those observed during the GTP-bound state (Supplementary Table S5). Altogether, our data indicates that (p)ppGpp induces a non-productive conformation of HflX, reducing the binding progression with the ribosomal subunit.

In the case of RbgA, all three tested nucleotides adhered to a two-step mechanism model, with *k*_app1_ increasing linearly with 50S concentration, while *k*_app2_ appeared hyperbolic (Supplementary Figure S5A, S5B). Further analysis to estimate the microscopic constants indicated that (p)ppGpp increased the dissociation rate constant *k*_-1_ by 3-5-fold as compared to GTP, whereas its association velocity *k*_1_ appeared largely unaffected (Supplementary Table S5, Supplementary Figure S5C, S5D). Interestingly, ppGpp drastically reduced the accommodation rate constant *k*_2_ similarly to HflX, while pppGpp did not. Altogether, our results indicate that (p)ppGpp can program RbgA to adopt different conformations that ultimately reduce their binding affinity for the ribosome (Supplementary Table S5).

For both RA-GTPases, the *K*_d_ of 50S binding is lower in the GTP-bound state compared to the (p)ppGpp-bound state (Supplementary Table S5). It appears that the main difference on a kinetic level, in agreement with our previous observations regarding the accommodation step, is that the binding of (p)ppGpp prevents the initiation of the slow-phase reaction (*k*_2_). (p)ppGpp would therefore prevent the stable association of the RA-GTPase with the ribosomal subunit. The affinity is further affected by the increased *k*_-1_ in the alarmone-bound states, which in addition to the lack of the second phase reaction while bound to (p)ppGpp, may lead to an increase in reversal reactions, enhancing the dissociation of the RA-GTPase from the ribosomal subunits. Altogether, the kinetic data are in accordance with the observation by western immunoblotting that RA-GTPases bind to target subunits more readily in the GTP-bound form but associate less readily in the presence of the stringent response alarmones (p)ppGpp. Specifically, (p)ppGpp appear to affect the forward reactions, consistent with inducing a non-productive conformation of the RA-GTPases. This could lead to diminished association of RA-GTPases to ribosomes at physiologically relevant alarmone concentrations, impairing ribosome maturation under stress.

### Association of the RA-GTPase Era to the 30S subunit decreases upon induction of the stringent response

Upon induction of the stringent response, cellular levels of (p)ppGpp increase, while concentration of GTP drops (38). Having observed decreased association of RA-GTPases to ribosomal subunits *in vitro*, we wished to examine the interaction under more physiologically relevant conditions. To investigate RA-GTPases interactions with the ribosome in the bacterial cell, we used an *era* deletion mutant in the community-acquired methicillin-resistant *S. aureus* (CA-MRSA) strain LAC* that was available to us. This strain has a growth defect (Figure 4A) and has an abnormal cellular ribosomal profile when compared to the wild-type, with an accumulation of 50S subunits and a loss of 70S ribosomes (Figure 4B, 4C) (10,27,42), suggesting that the absence of this GTPase is preventing mature ribosome formation and growth. In order to establish whether induction of the stringent response in bacterial cells leads to a decrease in the association of Era to the 30S subunit, the mutant was complemented with an anhydrotetracycline-inducible 6xHis-tagged version of *era*, yielding strain LAC* *Δera* iTET-*era*-His. Having confirmed that the His-tagged version of the protein is expressed and restores the growth defect observed in *era* mutant strains (Figure 4A), we grew cells to exponential phase and induced the stringent response with mupirocin, an antibiotic that inhibits isoleucyl tRNA synthetase and is known to activate the stringent response in *S. aureus* (43). Cells were lysed and applied to 10-40% sucrose gradients in ribosome dissociation buffer for subunit separation via isopycnic ultracentrifugation. The 30S pool was analysed for associated Era-His using *α*-His western immunoblotting (Figure 4D). Crude lysates sampled prior to loading on the sucrose gradients were also analysed to ensure equal loading and equal expression of Era-His between samples (Supplementary Figure S6). In agreement with the *in vitro* data, the relative association of Era-His to the ribosome decreased at least 4-fold upon induction of the stringent response (Figure 4D). Altogether this *in vitro* and *in vivo* data support a model in which the stringent response impairs 70S ribosome assembly by disrupting the association of RA-GTPases with the immature ribosomal subunits, thus preventing correct ribosome maturation.

**Figure 4.**
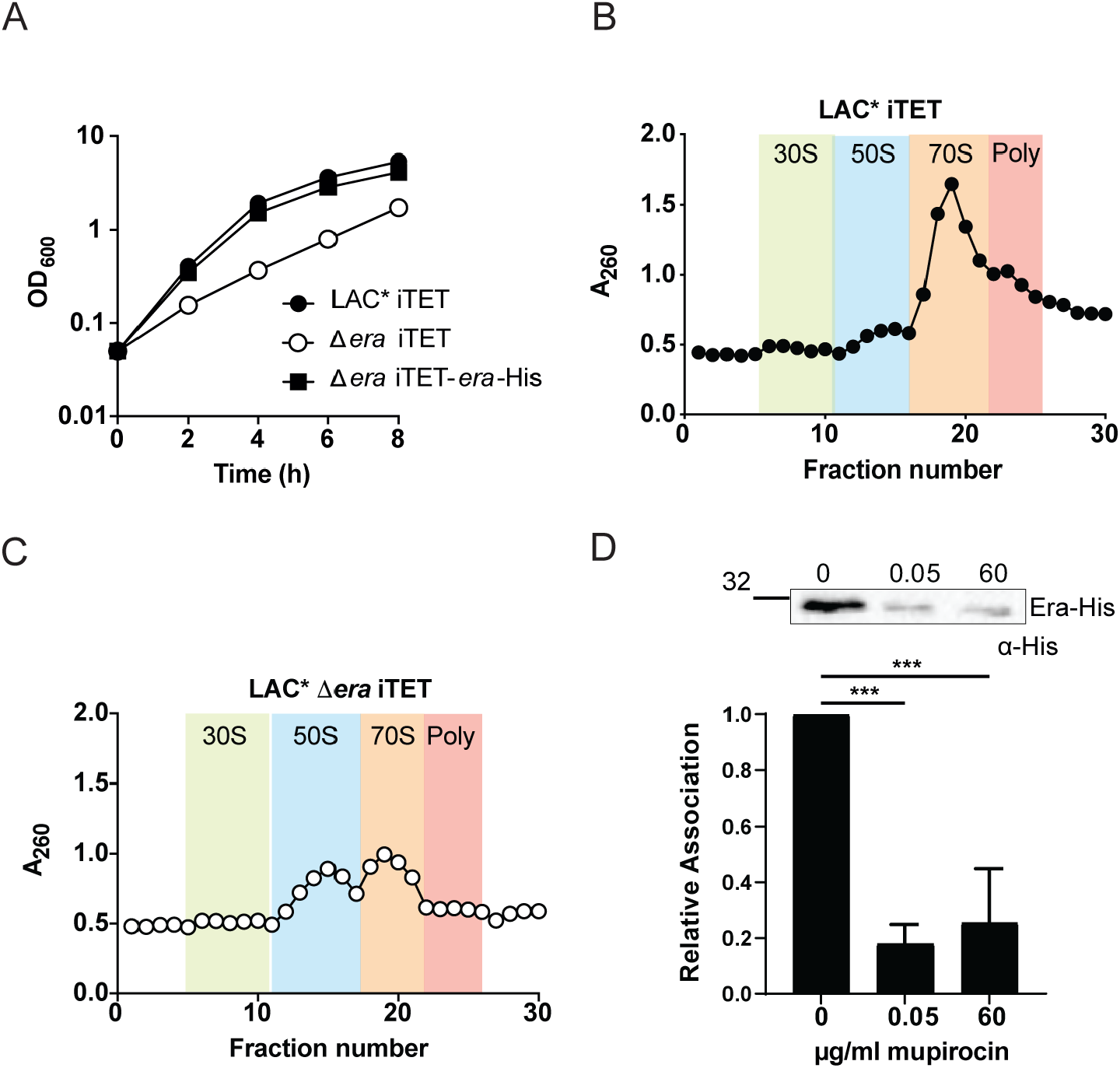
Association of Era to the 30S subunit is reduced under stringent conditions. A) Growth curve of *S. aureus* strains LAC* iTET, LAC* Δ*era* iTET and LAC* Δ*era* iTET-*era*-His. Overnight cultures were diluted to an OD_600_ of 0.05 and grown for 8 h in the presence of 100 ng/µl Atet. Experiments were carried out in triplicate, with error bars representing standard deviation. B, C) Ribosome profiles of the *S. aureus* strains B) LAC* iTET and C) LAC* Δ*era* iTET. RNA content was analysed at an absorbance of 260 nm. All experiments were performed in triplicate, with one representative profile included for each strain. Expected regions for 30S subunits (green), 50S subunits (blue), 70S ribosomes (orange) and polysomes (pink) are highlighted. D) Ribosome association of Era-His from strain LAC* Δ*era* iTET-*era*-His. Top: western immunoblot showing the association of Era-His to 30S ribosomes. LAC* Δ*era* iTET-*era*-His was grown to an OD_600_ of 0.5 in the presence of Atet. Cells were either left uninduced or grown in the presence of 0.05 or 60 µg/ml mupirocin for 30 mins to induce the stringent response. Ribosomal subunits were separated and the amount of Era-His associated was detected using HRP-conjugated α-His antibodies. Experiments were carried out in triplicate and one representative image is shown. Bottom: the mean signal intensities relative to the zero mupirocin sample of all repeats were plotted with error bars representing standard deviation. Statistical analysis was carried out using a one-way ANOVA followed by Tukey’s multiple comparisons test (*** *P* < 0.001).

### Crystallisation of RsgA in the apo and ppGpp-bound states

GTPases act as molecular switches, cycling between OFF (GDP-bound) and ON (GTP-bound) states. Structural studies of numerous GTPases have reported distinct conformations for both states, which are determined by the movement of the flexible switch I/G2 loop and the switch II/G3 loop (44). Often described as a loaded-spring mechanism, the conformational change occurs upon hydrolysis of GTP or the subsequent *γ*-phosphate release. Both switch I/G2 and switch II/G3 are responsible for coordinating the Mg^2+^ cofactor which interacts with the *γ*-phosphate of GTP via a conserved threonine residue in G2 and a glycine in G3. Upon hydrolysis of the *γ*-phosphate and P_i_ dissociation, the protein relaxes into the OFF conformation.

To look more at the mechanism of (p)ppGpp-mediated inhibition of RA-GTPases associating with ribosomal subunits, we solved the structures of RsgA in both the apo-(Figure 5A) and ppGpp-bound (Figure 5B) states by X-ray crystallography (Supplementary Table S3), in order to compare to already available GMPPNP- and GDP-bound structures. The 1.94 &Å structure of RsgA complexed with ppGpp reveals the presence of the nucleotide unambiguously represented in the electron density map (Supplementary Figure S7A), whereas the apo structure at 2.01 &Å is lacking any electron density in the nucleotide binding pocket. The overall structure of RsgA consists of three domains, the N-terminal OB-fold, the central GTPase domain and a C-terminal ZNF (Figure 5A). Both the OB-fold and ZNF domains are involved in nucleotide recognition (45,46), and target RsgA to the 30S ribosomal subunit where they contact major helices of the 16S rRNA (Figure 5C). The OB-fold is situated between h18 and h44, with the loop connecting β_1_ and β_2_ recognising the minor groove of h44 adjacent to the 30S acceptor site (4). The ZNF contacts the 30S head domain, making backbone contacts with h29 and h30, close to the interaction site of the P-site tRNA (4,47). In *E. coli* RsgA (YjeQ), the G-domain also contacts h44 by means of a clamp adjacent to the interaction site of h45 and h24. This clamping interaction is facilitated by the β_6,7_ hairpin and the switch I/G2 region (4), however this hairpin is lacking in *S. aureus* RsgA (Figure 5A, 5B), so it is likely that the G-domain interacts with h44 singly through the switch I/G2 region. The ppGpp ligand is bound in an elongated conformation, where the 3’ and 5’-phosphate moieties face away from each other (Supplementary Figure S7A). The guanosine-5’-diphosphate backbone interacts with the G-domain in an identical manner to the more well-characterised GMPPNP (Supplementary Figure S7B, S7C) (4,47), with the P-loop/G1 motif stabilising the *α, β*-diphosphate and the G4 motif specifically recognising the guanine nucleotide base. The 3’-diphosphate extends away from the core of the protein, towards the solvent and appears to be stabilised only by a long-range 5.5 &Å electrostatic interaction between the lone electron pair on the *ε*-phosphate of ppGpp and the basic lysine residue K116 (Supplementary Figure S7C). It is worth noting that in the GTP-bound ON state, the switch I/G2 and switch II/G3 flexible loops would aid in stabilising both the catalytic Mg^2+^ ion and *γ*-phosphate (4,47). In our structures there is no electron density corresponding to the Mg^2+^ and the switch I/G2 loop is unresolved, likely due to innate flexibility when not contacting a *γ*-phosphate. Additionally, the switch II/G3 loop does not appear to form hydrogen-bonds or electrostatic interactions with the ligand.

**Figure 5.**
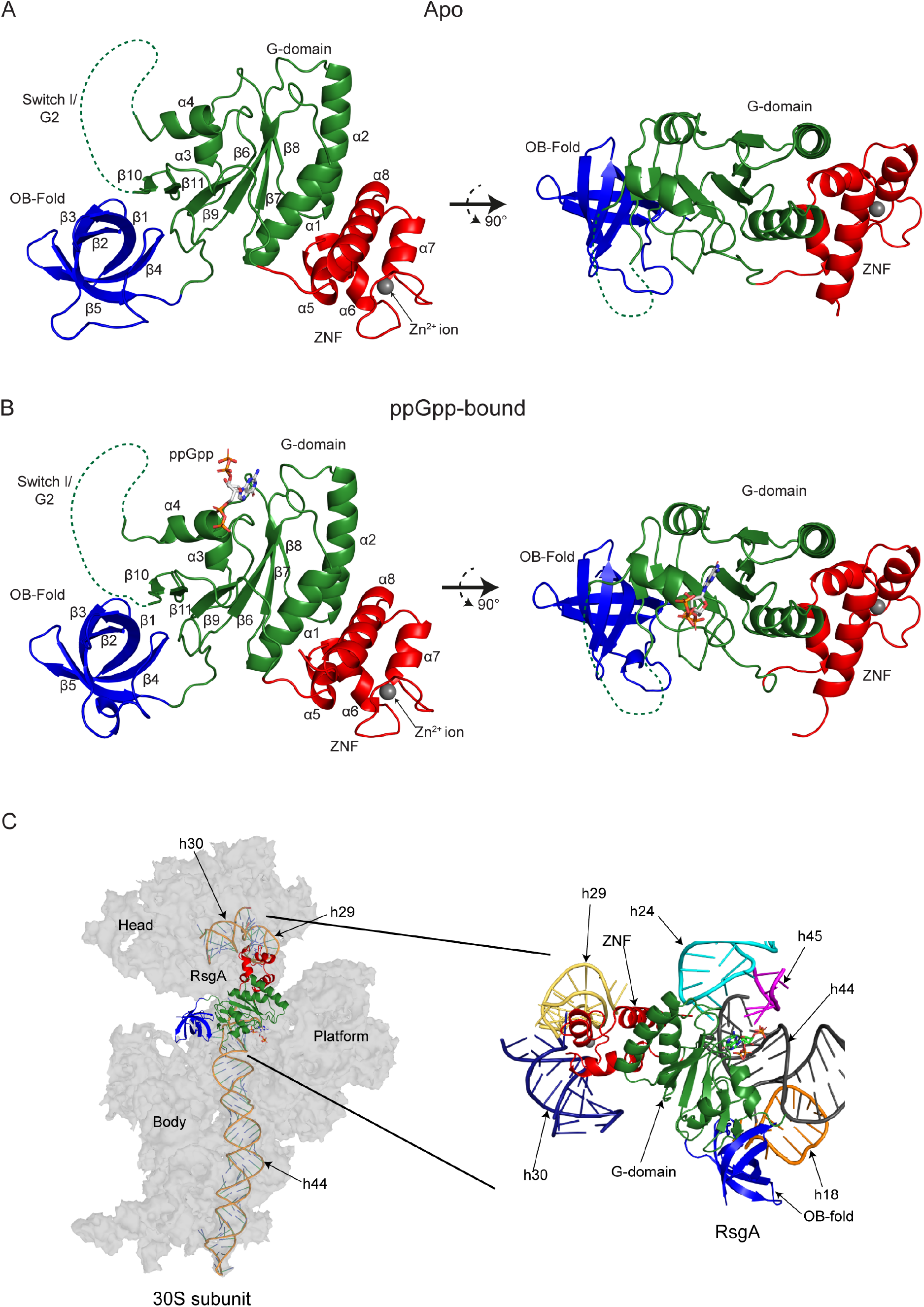
Structure of RsgA in the apo- and ppGpp-bound states. A, B) The crystal structures of RsgA in the apo state (A: PDB: 6ZJO) and bound to ppGpp (B: PDB: 6ZHL). The structures are coloured by domain, with the N-terminal OB-fold coloured blue, the central G-domain coloured green and the C-terminal Zn^2+^-finger (ZNF) domain coloured red. Structural details including α-helices, β-sheets, ligands and domains are labelled. The expected position of the switch I/G2 loops as determined by comparison with RsgA homologues in the GMPPNP-bound state are indicated using a dotted line, despite the lack of electron density surrounding this feature. For both A) and B), a 90° rotation around a horizontal axis is shown. C) The RsgA binding site on the 30S ribosomal subunit. RsgA-ppGpp (PDB: 6ZHL, this study) was overlaid onto the model of YjeQ-GMPPNP (PDB: 5UZ4, chain Z (47)) using C_α_ alignment, relative to the 30S ribosomal subunit (PDB: 5UZ4, chain A (47)). The RsgA model is shown as a cartoon representation, coloured by domain as above. The 30S subunit is shown in grey, with interacting rRNA helices shown as cartoon representations to highlight the RsgA recognition sites, as labelled. The bound ppGpp ligand is coloured by atom: carbon, grey; nitrogen, blue; oxygen, red; phosphorous, orange. Inset: a cartoon representation of the rRNA helices which constitute the RsgA binding site on the 30S subunit. Target rRNA helices are coloured as follows: h24, cyan; h18, orange; h29, yellow; h30, navy blue; h44, grey; h45, magenta.

### ppGpp-bound RsgA mimics the GDP-bound OFF-state conformation

For RsgA, a catalytic histidine residue is located within the switch I/G2 loop, two residues upstream of the conserved G2 threonine (4). Therefore, correct docking of this region upon binding to either GTP or the 16S rRNA is thought to be instrumental for GTPase activity. It has also been previously proposed by Pausch *et al*. (6) that for RbgA, the 3’-diphosphate of (p)ppGpp prevents the movement of switch I/G2 into the ON conformation necessary for GTP hydrolysis and ribosome binding, explaining why the GTPase is incapable of hydrolysing (p)ppGpp in a similar manner to GTP (6). In order to determine whether a similar steric inhibition is occurring for RsgA, we compared our apo and ppGpp-bound structures with available structures of RsgA homologues, namely *Aquifex aeolicus* YjeQ bound to GDP (PDB: 2YV5) and *E. coli* YjeQ complexed with both the 30S subunit and GMPPNP (PDB: 5UZ4 (47)) (Figure 6). Importantly, in both of these available structures, the switch I/G2 loops were partially resolved (Figure 6A, 6B). Despite a similar overall fold of the G-domain, the switch I/G2 loop in the GDP-bound structure appears to extend distally from the main body of the protein, far from the associated ligand (Figure 6A). Contrary to this, the GMPPNP-bound structure features a fully docked Switch I/G2 loop, positioned adjacent to the bound ligand and the binding site of the Mg^2+^ ion, although the Mg^2+^ ion itself is not resolved. Crucially, in this conformation, the docked switch I/G2 loop occupies the same space that the 3’-diphosphate moiety of ppGpp would (Figure 6B, 6D). Additionally, the switch II/G3 loop conformation differs between the GDP- and GMPPNP-bound structures, being extended towards the γ-phosphate of GMPPNP in the latter. When compared to our apo (Figure 6C) and ppGpp-bound (Figure 6D) structures, the switch II/G3 region appears highly similar to that of the GDP-bound structure, leading us to hypothesise that the switch I/G2 loop will also adopt a similar conformation to the GDP-bound state due to steric inhibition by ppGpp. This lack of docking of switch I/G2 would inhibit GTPase activity by preventing proper docking of the catalytic histidine within switch I (4), coordination of the Mg^2+^ cofactor by the G2 threonine (6), and subsequent interaction with the γ-phosphate of GTP.

**Figure 6.**
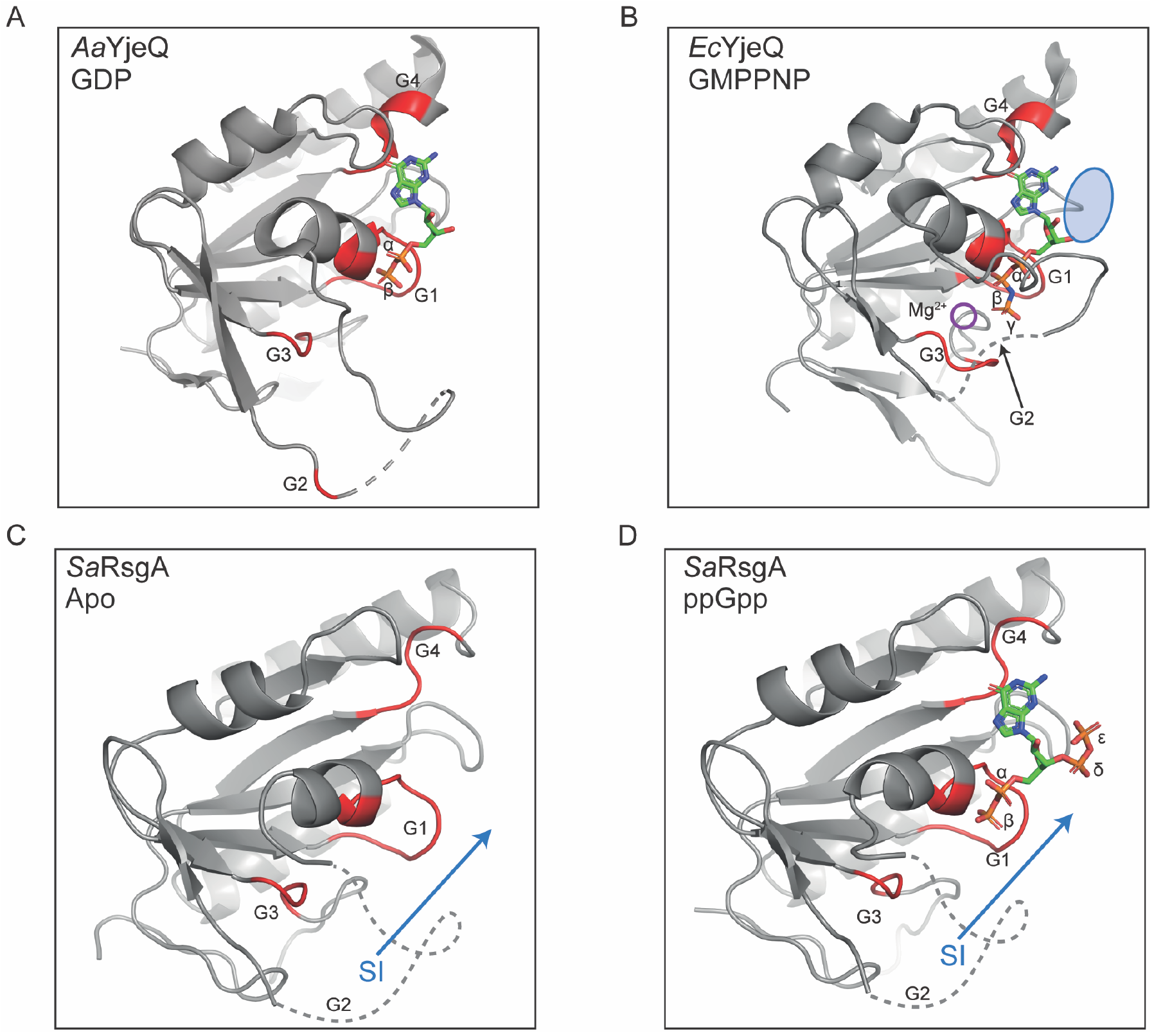
Comparison of the GTPase domains of RsgA and homologues in different nucleotide bound states. The G-domain conformation of A) *Aquifex aeolicus* RsgA (YjeQ) bound to GDP (PDB: 2YV5, chain A), B) *Escherichia coli* RsgA (YjeQ) bound to GMPPNP (PDB: 5UZ4, chain Z (47)), C) *Staphylococcus aureus* RsgA in the apo state (PDB: 6ZJO, chain A, this study) and D) *Staphylococcus aureus* RsgA bound to ppGpp (PDB: 6ZHL, chain A, this study). RsgA/YjeQ is shown as cartoon representations, coloured grey, with the G1, G2, G3 and G4 motifs coloured red where resolved. The hypothetical position of the switch I/G2 loop are represented by grey dashed lines as determined by comparison with the resolved region of the GDP-bound YjeQ, and the bound nucleotides are coloured by atom as follows: carbon, green; nitrogen, blue; oxygen, red; phosphorous, orange. Rearrangements of the switch I/G2 loop to facilitate entry into the ON state are shown by blue arrows. The binding site of the Mg^2+^ ion in the GMPPNP-bound conformation (B) is indicated by a purple circle, and the position of the δ, ε-phosphate of ppGpp is indicated relative to bound GMPPNP by a blue oval in B.

### Displacement of the G2 loop by (p)ppGpp inhibits RA-GTPase-ribosome interactions

The structure of RsgA in the GMPPNP-bound ON state has only ever been solved when associated with the 30S ribosomal subunit suggesting it is stabilised in this conformation (4,47). In order to assess the role of the switch I/G2 loop in ribosome association, we performed computational C_*α*_ alignments of both the available GDP-bound (PDB: 2YV5) and our ppGpp-bound structures with the GMPPNP-bound RsgA-30S ribosome complex (PDB: 5UZ4) (Figure 7A-C). It has previously been shown that each of the 3 domains of RsgA interact with rRNA to provide a stable docking interaction (Figure 5C) (4), and that for *E. coli* RsgA, the switch I/G2 loop and a *β*6, *β*7-hairpin clamp around h44, contacting the minor and major groove respectively (Figure 7A). However, when the GDP-bound OFF-state structure from *A. aeolicus* is superimposed in place of the GMPPNP structure, it appears that the switch I/G2 loop is positioned in such a way that would cause steric clashing between the phosphate backbone of h44 (Figure 7B). Likewise, the expected position of the switch I/G2 loop in the ppGpp-bound model would lead to similar steric clashing, with the 3′-diphosphate moiety of ppGpp preventing the switch I /G2 loop adopting the active conformation (Figure 7C). While it is important to stress that this modelling is performed using protein models and 30S subunits from separate organisms, this leads us to hypothesise that the misalignment of the switch I/G2 loop and subsequent steric clashing between the RA-GTPase and h44 of the 16S rRNA could be responsible for (p)ppGpp-mediated inhibition of RA-GTPase association to the ribosome. We suggest that this region is not directly responsible for promoting subunit docking, however that the switch I region instead forms electrostatic interactions with conformationally mature h44 and h45 rRNA following ribosome association, enabling positioning of the switch I/G2 loop in a catalytically active conformation when the mature rRNA conformation is reached. These interactions and the subsequent loop rearrangement may represent the slow stabilisation step (*k*_2_) observed in our stopped flow analysis (Figure 3).

**Figure 7.**
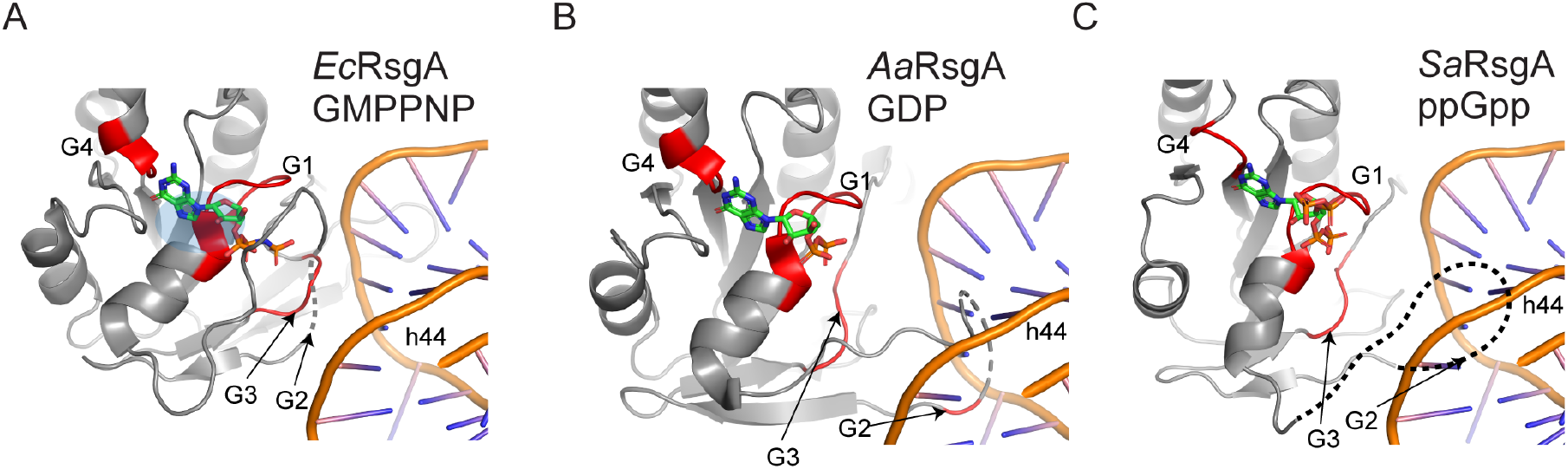
ppGpp-mediated inhibition of RA-GTPase association to ribosome subunits is facilitated by incorrect positioning of the switch I/G2 loop. A) *E. coli* RsgA (YjeQ) bound to GMPPNP (PDB: 5UZ4, chain Z) and chain A (16S rRNA) (47)) including a cropped view of the rRNA binding site on h44. For the full binding environment, see Figure 5. B, C) Docking of B) *A. aeolicus* RsgA (YjeQ) (PDB: 2YV5, chain A) bound to GDP and C) *S. aureus* RsgA bound to ppGpp (PDB: 6ZHL, chain A, this study) onto h44 of the 16S rRNA from PDB:5UZ4 (chain A) using C_α_ alignment of the G-domains. The RsgA/YjeQ is shown as a cartoon representation coloured grey, with the G1, G2, G3 and G4 motifs coloured red where visible. The bound nucleotides are coloured by atom as follows: carbon, green; nitrogen, blue; oxygen, red; phosphorous, orange. Suggested conformations of unresolved regions of the switch I/G2 loop are represented by a black dashed line.

## DISCUSSION

The stringent response is a multi-faceted stress coping mechanism, ubiquitously used throughout the Bacteria to cope with nutrient starvation conditions. Recent transcriptomics data has highlighted the diversity and complexity of this response, with 757 genes being differentially regulated within 5 minutes of (p)ppGpp induction (15). For Gram-positive bacteria, the regulation of transcription by (p)ppGpp is intricately linked to purine nucleotide levels, which are impacted in a number of ways (48). Upon induction of the stringent response, GTP/GDP and ATP levels decrease as they are utilised by (p)ppGpp synthetase enzymes (12). Furthermore, once produced (p)ppGpp directly inhibits a number of enzymes involved in the guanylate and adenylate synthesis pathways, further reducing GTP/GDP levels (38,49). All of this results in a shift from high GTP/GDP and low (p)ppGpp levels in fast growing cells, to low GTP/GDP and high (p)ppGpp in nutritionally starved cells. For *S. aureus*, the impacts of this are wide-reaching, affecting transcription initiation (39), enzyme activities (50) and, as we show here, the regulation of the activity of RA-GTPases by tuning their capacity to interact with ribosomal subunits.

In the present work, we examine the nucleotide binding preferences of RA-GTPases, and the consequences of this binding on regulating the interactions of RA-GTPases with the ribosome. Cycling between the GTP-bound ON and GDP-bound OFF states is critically important for RA-GTPases, as it enables these proteins to act as molecular checkpoints of ribosome assembly. Here we show that RA-GTPases bind to guanosine nucleotides competitively and with differing affinities, with GDP and ppGpp binding with up to 6-times greater affinity than their 5’ trinucleotide-containing counterparts GTP and pppGpp (Supplementary Table S4). The consequence of differing nucleotide-bound states for interactions with ribosomal subunits are significant. We observe that GTP binding is required to promote RA-GTPase/ribosome interactions (Figure 2 & 3). Indeed, the binding of apo RbgA and HflX to the 50S subunit was almost undetectable by stopped-flow fluorescence (Figure 3B, 3C), although Era and HflX did demonstrate background binding to the 30S and 50S subunits respectively by western immunoblotting. A cryo-electron micrograph (cryo-EM) structure of Era binding to the 30S subunit has previously been solved (40), demonstrating that this GTPase can bind in the apo form. Upon induction of the stringent response, levels of (p)ppGpp in the cell rise, swiftly becoming the dominant guanosine nucleotide in the cell (12,51), causing (p)ppGpp to out-compete GTP for occupancy of the nucleotide binding site (Figure 1C, Supplementary Figure S2D-F), and resulting in reduced association of RA-GTPases to their target ribosomal subunit and thus reduced production of mature 70S ribosomes (Figure 2, 3 & 4). It has been previously shown that, opposite to our observations regarding ribosome assembly factors, ppGpp binding enhances the affinity of the (p)ppGpp-binding RA-GTPase ObgE to the 50S subunit (52). This may reflect the proposed role of ObgE as a 50S based late-stage anti-association factor (52) which would benefit from enhanced affinity for the 50S in the ppGpp-bound state to prevent subunit joining and 70S formation. Unfortunately, we were unable to purify enough ObgE to compare using our system. Structural studies indicate that ppGpp-bound RsgA and RbgA most likely mimic the GDP-bound OFF state (Figure 5 & 6)(6), with (p)ppGpp inhibiting both GTPase activity and ribosome association by displacement of the switch I/G2 loop into an OFF-state conformation not compatible with stable ribosome subunit interaction (Figure 7). Given the reaction scheme determined by stopped-flow fluorescence, it is possible that the slower stabilisation step (*k*_2_) observed when HflX is in the ppGpp-bound state compared to the GTP-bound state could be due to improper loop docking following association of the RNA-binding domain(s) with the ribosome, leading to dissociation.

Altogether, our data favour a model (Figure 8) whereby in unstressed growing cells, GTP is the predominant nucleotide and induces the RA-GTPase ON-state conformation. Binding of the enzymes to each individual ribosomal subunit follows in order to promote a processing event. Following this, GTP is hydrolysed to GDP, inducing formation of the OFF state and subsequent dissociation. Upon cell starvation, the concentration of (p)ppGpp in the cell rises sharply, where it can out-compete GTP for binding to the RA-GTPases. The increase in (p)ppGpp not only inhibits the GTPase activity, but also negatively impacts the stability of RA-GTPase-ribosome interactions, reducing biogenesis and slowing growth. With its high concentration and affinity, (p)ppGpp could remain bound to RA-GTPases, preventing further ribosome biogenesis during low-energy conditions, yet preserving a pool of enzymes ready for rapid resumption of growth upon restoration of the proliferative state.

**Figure 8.**
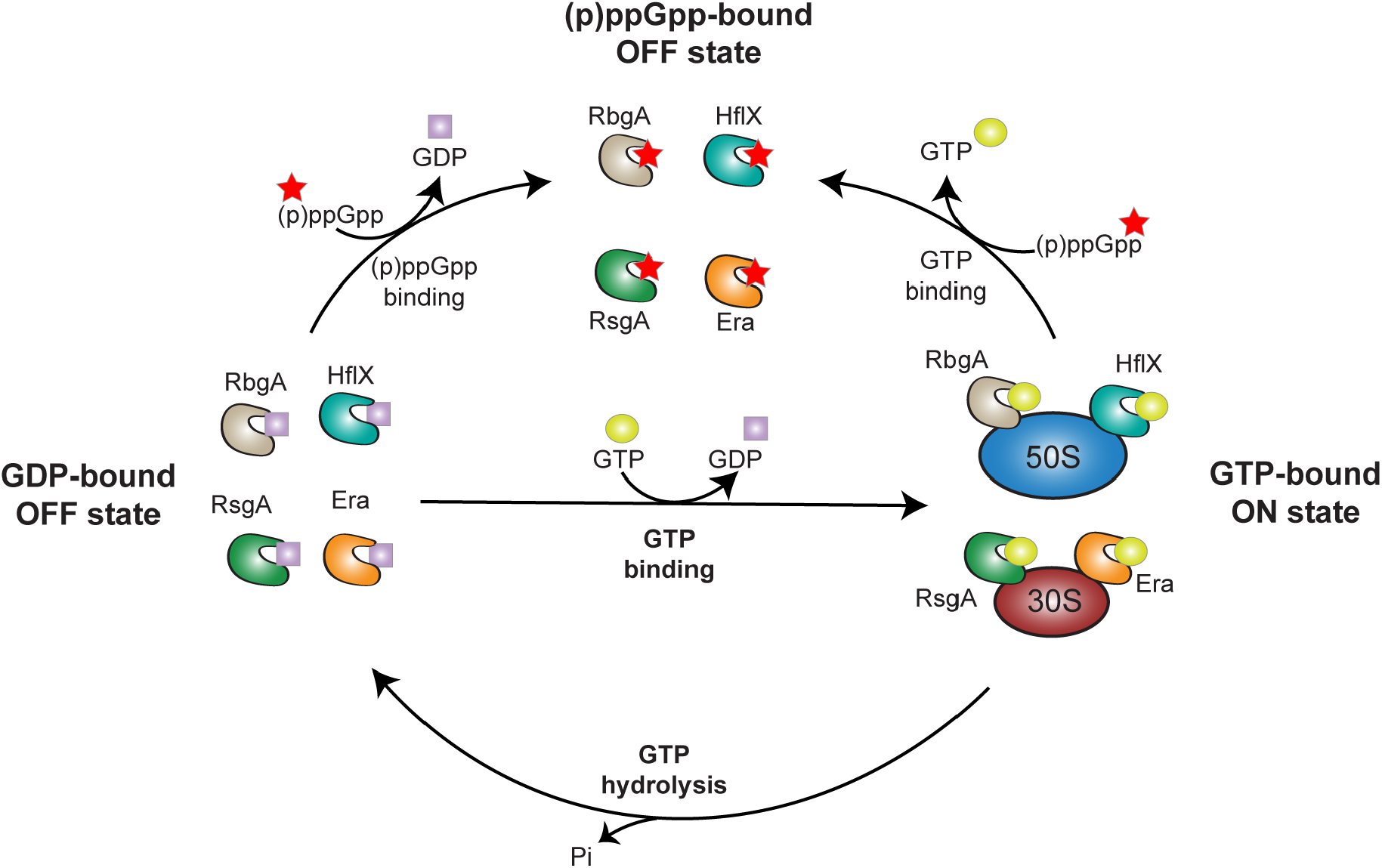
Model of the control of ribosome maturation by (p)ppGpp and RA-GTPases. Under proliferative conditions, GTP binds to RA-GTPases, enabling association to the immature ribosome subunits and subsequent maturation, at which point GTP is hydrolysed to GDP and the RA-GTPase dissociates from the ribosome. Under stringent conditions when cellular (p)ppGpp concentrations rise and GTP concentrations fall (38,64), (p)ppGpp can outcompete GTP for RA-GTPase binding. This inhibits association of RA-GTPases to the ribosome subunits and negatively impacts ribosome biogenesis.

All of the above analyses suggest that both pppGpp and ppGpp mimic the GDP-bound OFF conformation, an assertion that we support by solving the crystal structure of RsgA in the apo and ppGpp-bound states (Figure 5 & 6). These structures are in line with the OFF-state conformations observed by Pausch *et al* for RbgA in complex with (p)ppGpp (6). Similar to our RsgA-ppGpp structure, the diphosphate moieties of ppGpp bound by RbgA are in the elongated conformation (6), where the 3’ and 5’-phosphate moieties face away from each other. This configuration is not consistent among all (p)ppGpp-binding proteins or even among RA-GTPases. For example, the *E. coli* RA-GTPases BipA and ObgE bind to ppGpp in a ring-like conformation (53-55), in which the 3’ and 5’ phosphate moieties point towards each other. While no structural reasoning for this difference in conformation is known, aside from to extend the breadth of responses controlled by (p)ppGpp, it has been suggested that proteins which bind (p)ppGpp in the ring-like conformation have 10-fold lower inhibitory constants and dissociation constants than those which bind in the elongated conformation (56,57). This could potentially influence the temporal or energetic threshold during the stringent response where a certain protein becomes inhibited, based on decreasing concentrations of GTP and increasing concentrations of (p)ppGpp (38).

Ribosomal rRNA production and biogenesis are not the only aspects of protein synthesis that (p)ppGpp regulates, given its ability to bind to the bacterial initiation factor 2 (IF2), elongation factor Tu (EF-Tu), elongation factor G (EFG), elongation factor Ts (EFTs) and release factor 3 (RF3) (58-62). In each case, competitive binding of (p)ppGpp to these GTPases results in an inhibition of activity and reduction of the elongation cycle. Unlike RA-GTPases involved in subunit maturation, both IF2 and EFG bind to GTP, GDP and (p)ppGpp with similar affinity (59,60,63), albeit with EFG demonstrating an overall lower affinity. Furthermore, IF2 binding to (p)ppGpp within the 30S pre-initiation complex alters the mRNA binding preference, enabling permissive translation of certain mRNAs such as *tufA* encoding EF-Tu (58), which may fine-tune the proteome on a translational level to better enable survival of nutrient deprivation.

With this work we have used complementary techniques to demonstrate that (p)ppGpp prevents stable association of RA-GTPases to the ribosome, both *in vitro* and within the bacterial cell. This is achieved by these enzymes having a stronger affinity for ppGpp over GTP, with ppGpp interactions holding these enzymes in an OFF-state conformation. Consequently, this imparts delays to 70S ribosome assembly, which in turn contributes to the growth defects that are observed upon induction of the stringent response. Altogether, we highlight RA-GTPases-(p)ppGpp interactions as important regulators of stringent response-mediated growth control.

## DATA AVAILABILITY

The coordinates and electron density maps of RsgA-apo and RsgA-ppGpp have been deposited in the Protein Data Bank in Europe (PDBe) (https://www.ebi.ac.uk/pdbe/node/1) under accession codes 6ZJO and 6ZHL respectively.

## SUPPLEMENTARY DATA

Supplementary data are available at NAR Online.

