## supplemental tables and figures for "(p)ppGpp inhibits 70S ribosome formation in *Staphylococcus aureus* by impeding GTPase-ribosome interactions"

**SUPPLEMENTARY INFORMATIONSUPPLEMENTARY TABLES**

**Table S1. Bacterial strains used in this study**

| Strain | Relevant features | Reference |
|---|---|---|
| | ***Escherichia coli* strains** | |
| XL1-Blue | Cloning strain: TetR | Stratagene |
| BL21(DE3) | Strain used for protein expression | Novagen |
| RMC0147 | pET28b in XL1-Blue: KanR | Novagen |
| RMC0169 | pET28b-*gppA* in BL21(DE3): KanR | (1) |
| RMC0178 | pET21a-*relseq* in BL21(DE3): CarbR | (2) |
| RMC0399 | pET28b-*rsgA* in BL21 (DE3): KanR | (1) |
| RMC0401 | pET28b-*rbgA* in BL21 (DE3): KanR | (1) |
| RMC0402 | pET28b-*era* in BL21 (DE3): KanR | (1) |
| RMC0403 | pET28b-*hflX* in BL21 (DE3): KanR | (1) |
| RMC0531 | pCN55iTET in XL1-Blue: CarbR | (3) |
| RMC1323 | pCN55iTET-*era*-his in XL1-Blue: CarbR | This study |
| | ***Staphylococcus aureus* strains** | |
| SEJ1 | RN4220 *spa;* protein A negative derivative of RN4220; ANG314 | (4) |
| LAC* | LAC*: Erm sensitive CA-MRSA LAC strain (AH1263) | (5) |
| RMC0562 | LAC* pCN55iTET: SpecR | (3) |
| RMC0650 | LAC* Δ*era*: TetR | (3) |
| RMC0813 | LAC* Δ*era* pCN55iTET: TetR, SpecR | (3) |
| RMC1690 | LAC* Δ*era* pCN55iTET-*era*-his: TetR, SpecR | This study |

Antibiotics were used at the following concentrations: for *E. coli* cultures: Kanamycin (KanR) 30 µg/ml, Carbenicillin (CarbR) 50 µg/ml; for *S. aureus* cultures: Tetracycline (TetR) 2 µg/ml, Spectinomycin (SpecR) 250 µg/ml; IPTG at 1 mM, and anhydrotetracycline (Atet) at 100 ng/ml.

**Table S2. Primers used in this study**

| Number | Name | Sequence |
|---|---|---|
| RMC068 | F-NdeI-Era | CCCCATATGACAGAACATAAATCAGCATTTGT |
| RMC069 | R-BamHI-Era | CCCGGATCCTTAATCTTGGTCTTCAACATAACC |
| RMC157 | F-KpnI-Era | GGGGGGTACCTCAGGAAAGGATTTAGAATAAATG |
| RMC536 | R-SacI-Era-His | AAAGAGCTCTTAGTGATGGTGATGGTGATGACCATCTTGGTCTTCAACATAACC |

Restriction sites in primer sequences are underlined

**Table S3. Crystallographic data and refinement statistics**

| | RsgA Apo | RsgA-ppGpp |
|---|---|---|
| **Crystal data** | | |
| Space Group | $P\ 1\ 2_1\ 1$ | $P\ 2_1\ 2_1\ 2_1$ |
| Unit Cell Dimensions (a, b, c (Å)) | 54.67 93.53 68.18 | 50.47 66.93 114.12 |
| Unit Cell Dimensions (α, β, γ (°)) | 90.00 90.72 90.00 | 90.00 90.00 90.00 |
| **Data Collection** | | |
| Wavelength (Å) | 0.97949 | 0.97949 |
| Resolution (Å) | 47.20-2.01 (2.06-2.01) | 57.73-1.94 (1.99-1.94) |
| Reflections (measure d/unique) | 306,023 | 375,925 |
| $R_{meas}$ (%) | 0.149 (0.895) | 0.165 (0.990) |
| $R_{p.i.m.}$ (%) | 0.057 (0.336) | 0.063 (0.381) |
| $\langle I/\sigma I \rangle$ | 8.4 (1.8) | 11.0 (2.5) |
| Multiplicity | 6.8 (7.0) | 12.8 (12.7) |
| Completeness (%) | 98.2 (97.0) | 99.9 (99.5) |
| **Refinement Statistics** | | |
| $R_{work}/R_{free}$ (%) | 22.41/27.83 | 21.63/26.22 |
| Average B factor (Å$^2$) protein | 38.423 | 29.398 |
| Average B factor (Å$^2$) solvent | 42.832 | 34.955 |
| Rmsd bond lengths (Å) | 0.0083 | 0.0096 |
| Rmsd bond angle (°) | 1.5890 | 1.8108 |
| Protein residues | 533 | 268 |
| Water molecules | 212 | 81 |

| | | |
|---|---|---|
| Ions | 5 | 1 |
| Ramachandran (Favoured/Generous/Disallowed) | 494/27/4 | 255/7/1 |

Outer shell data in parenthesis. $R_{work} = \frac{\Sigma||F_{obs}|-|F_{calc}||}{\Sigma|F_{obs}|}$, where $F_{obs}$ and $F_{calc}$ are the observed and calculated factorial amplitudes of the structure respectively. $R_{free}$ is calculated as above, except for a random subsection of data that was withheld from refinement. Ramachandran plot calculated within *Coot*. B factors calculated using Baverage within the CCP4 suite. Refinement statistics were read from the output log following crystallographic refinement via RefMac5 within the CCP4 suite (6,7).

**Table S4. Binding affinities**

| | Era | | RbgA | | RsgA | | HflX | |
|---|---|---|---|---|---|---|---|---|
| | $K_d$ (μM) | Bmax | $K_d$ (μM) | Bmax | $K_d$ (μM) | Bmax | $K_d$ (μM) | Bmax |
| **GDP** | 4.94 ± 0.72 | 0.54 ± 0.02 | 6.07 ± 1.05 | 0.52 ± 0.02 | 1.83 ± 0.16 | 0.92 ± 0.02 | 4.91 ± 0.70 | 0.68 ± 0.02 |
| **ppGpp** | 4.21 ± 0.55 | 0.42 ± 0.01 | 2.86 ± 0.40 | 0.56 ± 0.02 | 2.17 ± 0.20 | 0.85 ± 0.02 | 3.37 ± 0.44 | 0.62 ± 0.02 |
| **GTP** | 11.50 ± 1.61 | 0.32 ± 0.01 | 18.48 ± 5.35 | 0.40 ± 0.04 | 3.56 ± 0.41 | 0.77 ± 0.02 | ND | 0.65 ± 0.34 |
| **pppGpp** | 13.87 ± 4.71 | 0.20 ± 0.02 | 13.76 ± 4.04 | 0.41 ± 0.03 | 10.06 ± 2.16 | 0.43 ± 0.03 | ND | 0.48 ± 0.22 |

**Table S5. Association ($k_1$, $k_2$) and dissociation ($k_{-1}$, $k_{-2}$) rate constants and approximate dissociation constant ($K_d$) of HflX and RbgA binding to 50S ribosomes in various nucleotide-bound states**

| | | $k_1$ ($s^{-1}$) | $k_{-1}$ ($s^{-1}$) | $k_2$ ($s^{-1}$) | $k_{-2}$ ($s^{-1}$) | $K_d$ (μM) |
|---|---|---|---|---|---|---|
| **HflX** | **GTP** | 20.7 ± 1.7 | 3.9 ± 0.5 | 1.6 ± 0.4 | 0.7 ± 0.3 | 0.06 ± 0.03 |
| | **ppGpp** | 8.9 ± 3.3 | 7.0 ± 1 | ~0 ± 0.5 | 1.1 ± 0.2 | 1.2 ± 0.9 |
| **RbgA** | **GTP** | 13.3 ± 0.9 | 1.3 ± 0.4 | 0.14 ± 0.7 | 0.6 ± 0.7 | 0.08 ± 0.08 |
| | **ppGpp** | 15.7 ± 6.7 | 3.6 ± 3.4 | ~0 ± 1.3 | 0.6 ± 1.2 | 0.3 ± 0.6 |
| | **pppGpp** | 12.6 ± 2.6 | 5.4 ± 1.3 | 0.4 ± 4.4 | 0.5 ± 4.4 | 0.2 ± 2 |

Negative rate values for $k_2$ approximated to ~0 $s^{-1}$. HflX complexed with pppGpp did not obey a two-step model for interaction and appeared rate-limited by an isomerization step at 5 $s^{-1}$ (Figure 3D). Error values shown represent the standard error of the two-stage analysis.

### SUPPLEMENTARY FIGURES

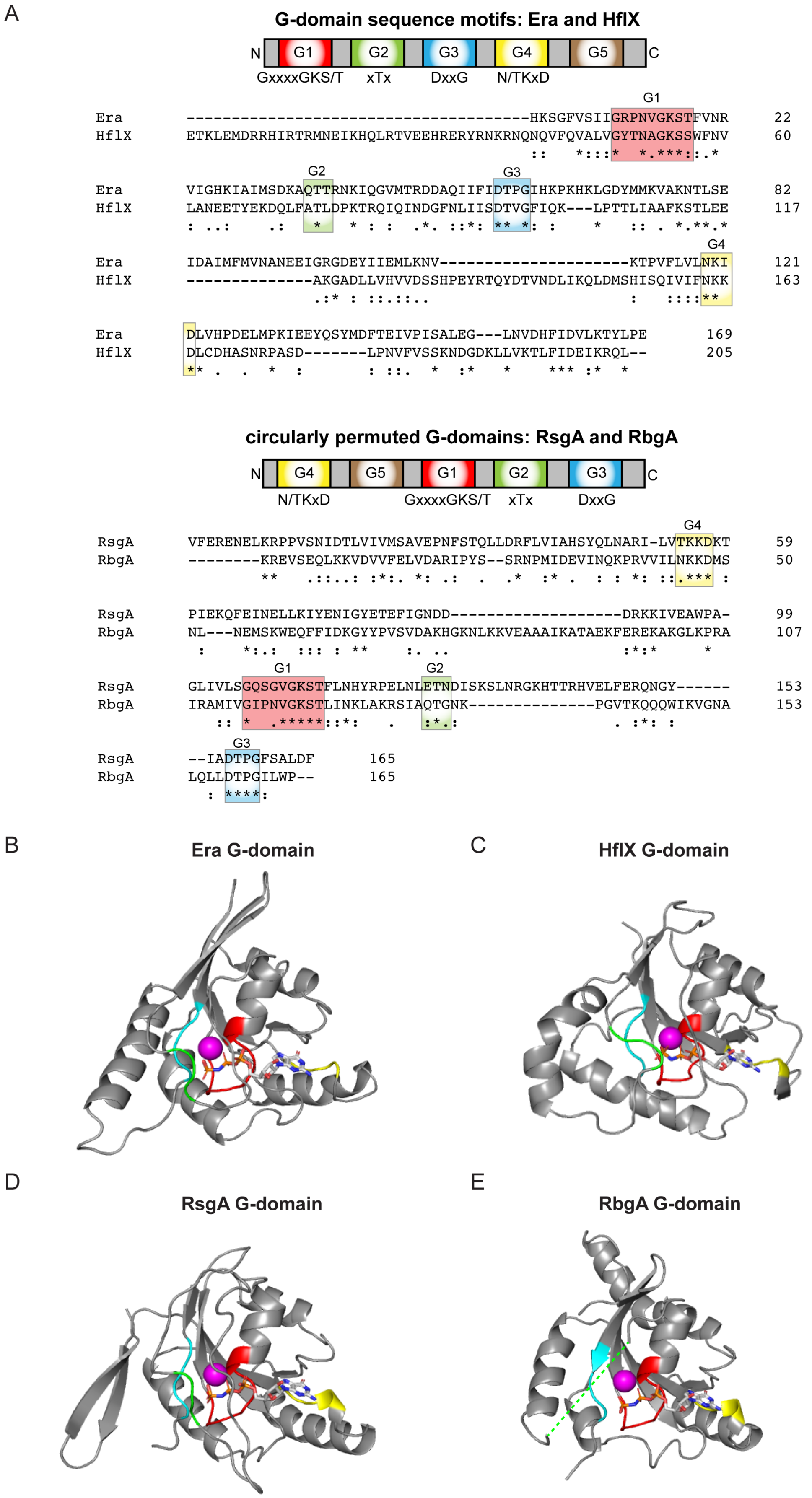

**Figure S1. Comparison of the GTPase domains of Era, HflX, RsgA and RbgA.** A) Domain structure and sequence alignments of the GTPase domains of *S. aureus* Era and HflX (top) and RsgA and RbgA (bottom). The conserved functional motifs G1-4 are highlighted as follows: G1, red; G2, green; G3, blue and G4, yellow. The G5 motifs (brown) are not shown in the alignment due to low sequence conservation. Note the circular permutation of the RsgA and RbgA GTPase domains results in the G4 and G5 motifs being located N-terminal of the G1, 2 and 3 motifs. Sequence alignments were generated using Clustal Omega (8). B-E) Structures of the GTPase domains of B) *Aquifex aeolicus* Era (adapted from PDB: 3R9W), C) *E. coli* HflX (adapted from PDB: 5ADY), D) *B. subtilis* YloQ (an RsgA homologue, adapted from PDB: 5NO3), and E) *B. subtilis* YlqF (an RbgA homologue, adapted from PDB: 1PUJ), in the GMPPNP-bound states. Functional motifs G1 (red), G2 (green), G3 (blue) and G4 (yellow) are highlighted. The bound GMPPNP ligand is coloured by atom and the bound $Mg^{2+}$ ion is shown in magenta. The switch I (G2) region of RbgA was unresolved in the structure, therefore the rough expected location of switch 1 is marked by a green dashed line.

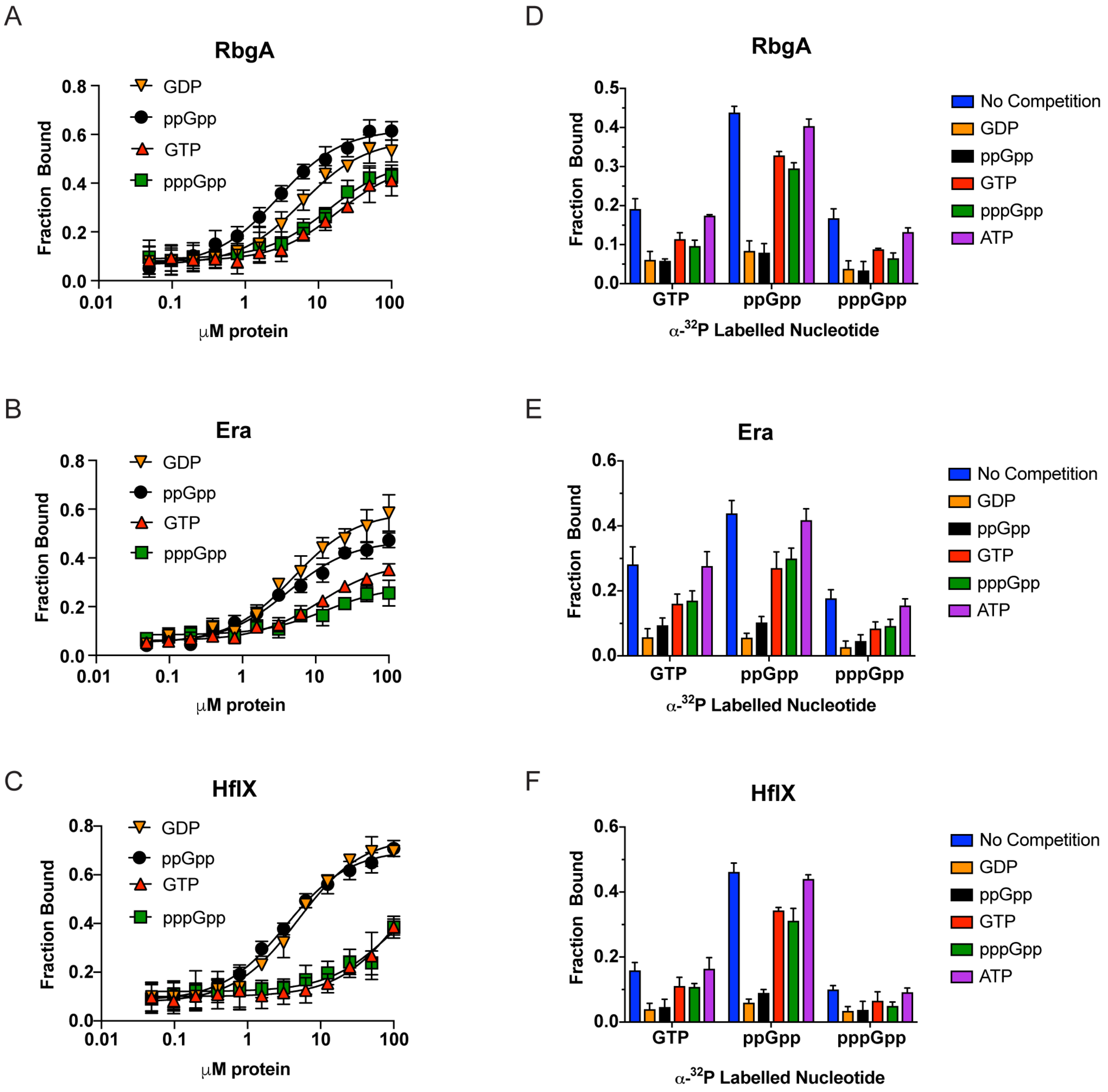


**Figure S2. Examination of the binding of GDP, ppGpp, GTP and pppGpp to RA-GTPases by DRaCALA.** A – C) Determination of binding affinities and $K_d$ values for $^{32}$P-labelled nucleotides to purified recombinant A) RbgA, B) Era and C) HflX. $K_d$ values were determined from the binding curves as previously described (9). D – F) DRaCALA binding assay of recombinant D) RbgA, E) Era and F) HflX binding to $^{32}$P-labelled GTP, ppGpp and pppGpp in the presence and absence of 100 μM cold competitors (GTP, GDP, ppGpp, pppGpp and ATP). All experiments were carried out in triplicate, with error bars representing standard deviation.

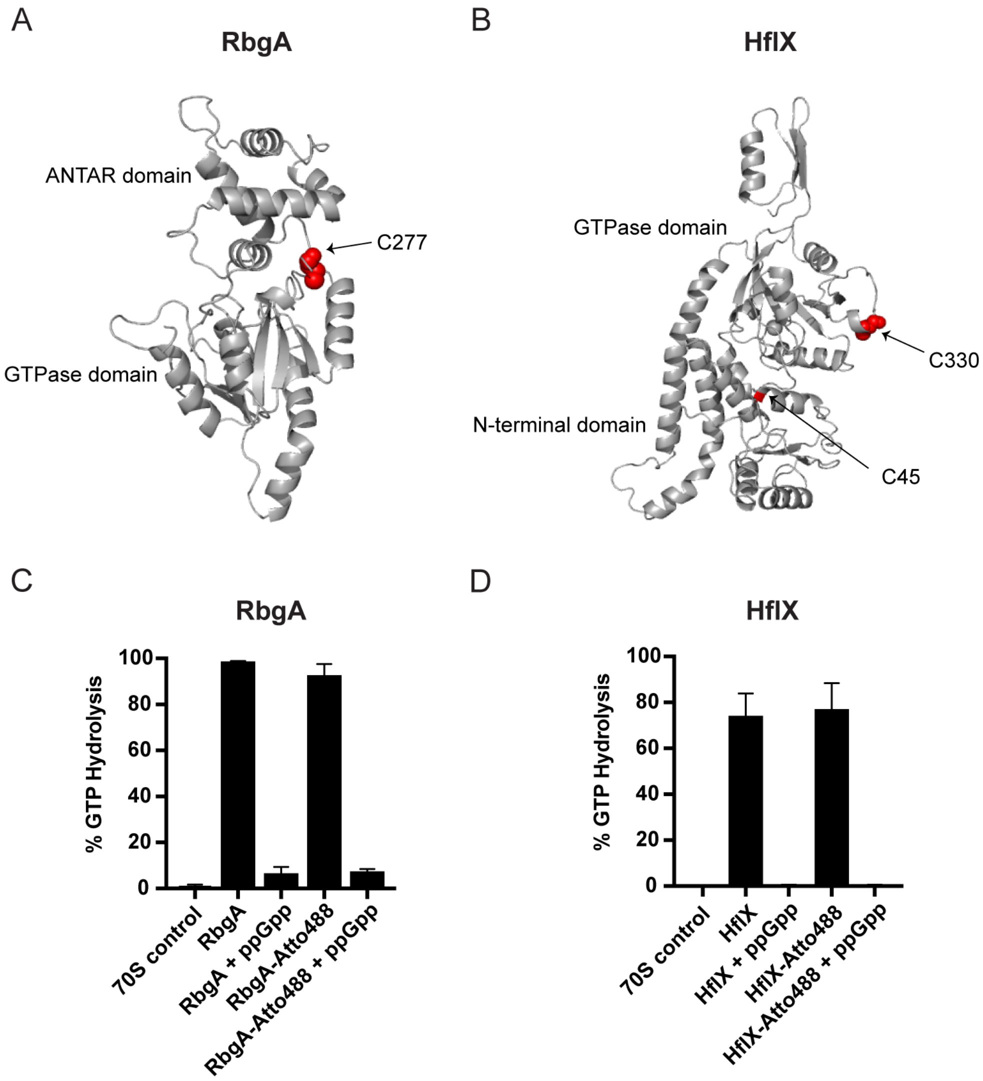


**Figure S3. Labelling sites and GTPase activity of Atto488-labelled proteins.** A, B) Predicted full-length structures of A) RbgA and B) HflX. Cysteine residues are shown in red, and protein domains are indicated. Cysteine residues amenable for labelling with Atto488-maleimide are shown as spheres. Structures were predicted through homology modelling (SWISS MODEL server) (10), using template PDBs 1PUJ (chain A) and 5ADY respectively. C, D) The GTPase activity of 0.1 μM wild-type and Atto488-labelled C) RbgA and D) HflX, in the presence and absence of 100 μM ppGpp. Reaction mixtures containing 1 μM GTP spiked with $^{32}$P-labelled GTP, 0.1 μM RA-GTPase and 0.1 μM 70S ribosomes to stimulate GTPase activity were incubated at 37°C for 60 minutes. Control reactions lack the RA-GTPases. Enzymatic activity was monitored by TLC and quantified using ImageQuant. Experiments were performed in triplicate, and error bars represent standard deviations.

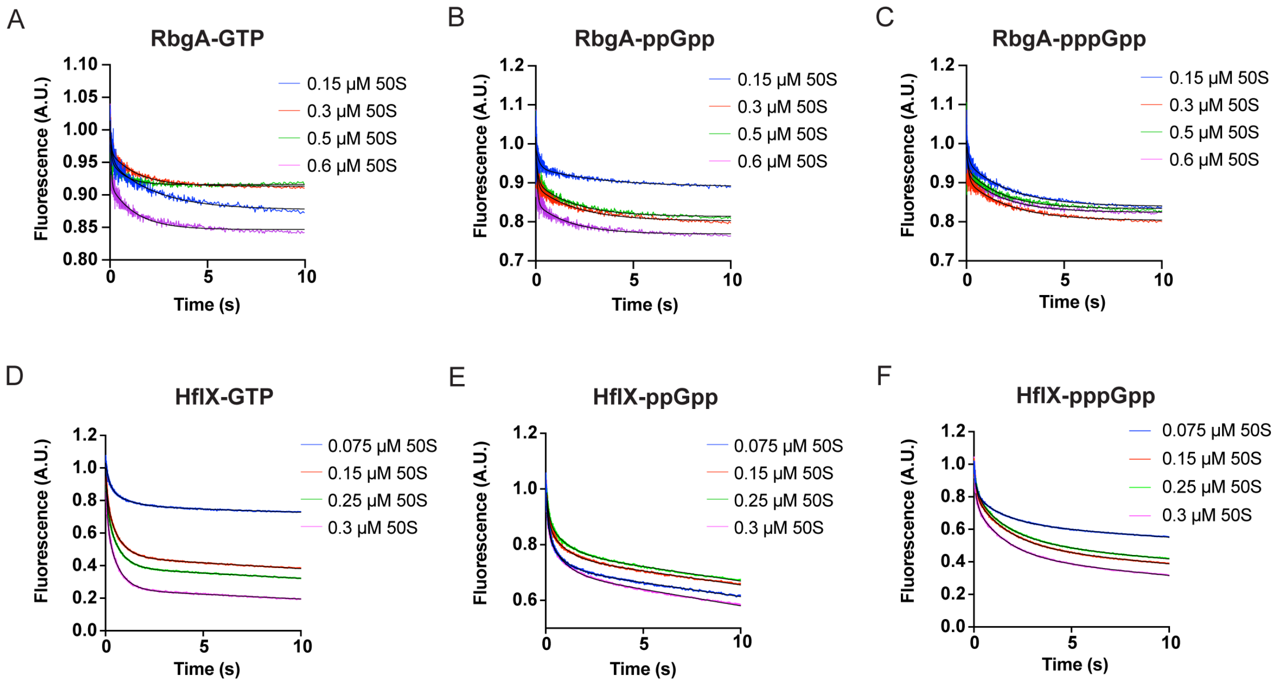


**Figure S4. Stopped-flow kinetic parameters of RA-GTPases associating to the 50S ribosome.** A – C) 0.075 μM Atto488-labelled RbgA was rapidly mixed with increasing molar excesses of 50S ribosomal subunits in the presence of A) GTP, B) ppGpp or C) pppGpp. D – F) 0.05 μM HflX-Atto488 was mixed with increasing molar excesses of 50S in the presence of D) GTP, E) ppGpp and F) pppGpp. Sampling was carried out in an exponential manner over a 10 second period, and resulting traces were analysed by nonlinear regression using two exponential terms, shown as a solid black line. Each trace is the average of at least 5 replicates.

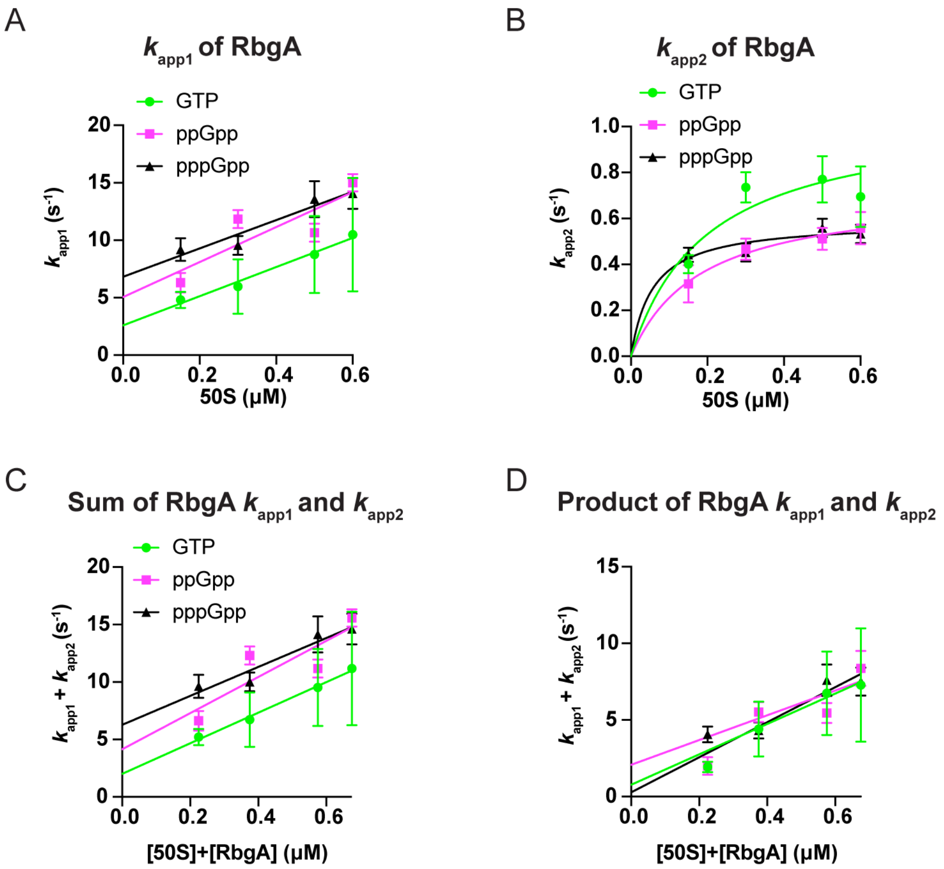


**Figure S5. Sum and product analyses of apparent rates during RbgA association to the 50S subunit.** A-D) 0.075 µM RbgA-Atto488 was mixed with increasing titrations of 50S ribosomal subunits at molar excess over fluorescently labelled protein, in the presence of 20 µM GTP, ppGpp or pppGpp. The resultant traces (Supplementary Figure S4) were analysed by nonlinear regression using two exponential terms. The sum (C) and product (D) of apparent rates ($k_{app1}$ (A), $k_{app2}$ (B)) were plotted as a function of the total concentration of the 50S subunit and RbgA protein and the dissociation constant ($K_d$) calculated as specified in the Methods section. Error bars represent the standard deviation of the apparent rates of 4 or more individual traces (A, B) or the standard error of the two-step analysis (C, D).

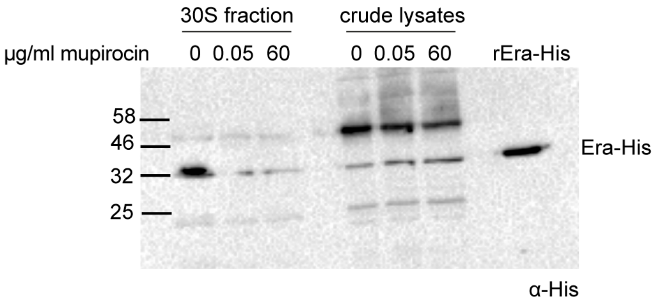


**Figure S6. Association of Era to the 30S subunit is reduced under stringent conditions.** Western immunoblot showing the association of Era-His to 30S ribosomes. LAC* Δ*era* iTET-*era*-His was grown to an $OD_{600}$ of 0.5 in the presence of Atet. Cells were either left uninduced or grown in the presence of 0.05 or 60 µg/ml mupirocin for 30 mins to induce the stringent response. Ribosomal subunits were separated and the amount of Era-His associated to the 30S fraction (left) or present in crude lysates prior to loading (right) was detected using HRP-conjugated α-His antibodies. Recombinant Era-His protein was also loaded as a control for protein size and identification (last lane). Experiments were carried out in triplicate and one representative image is shown. Ladder sizes are indicated on the left.

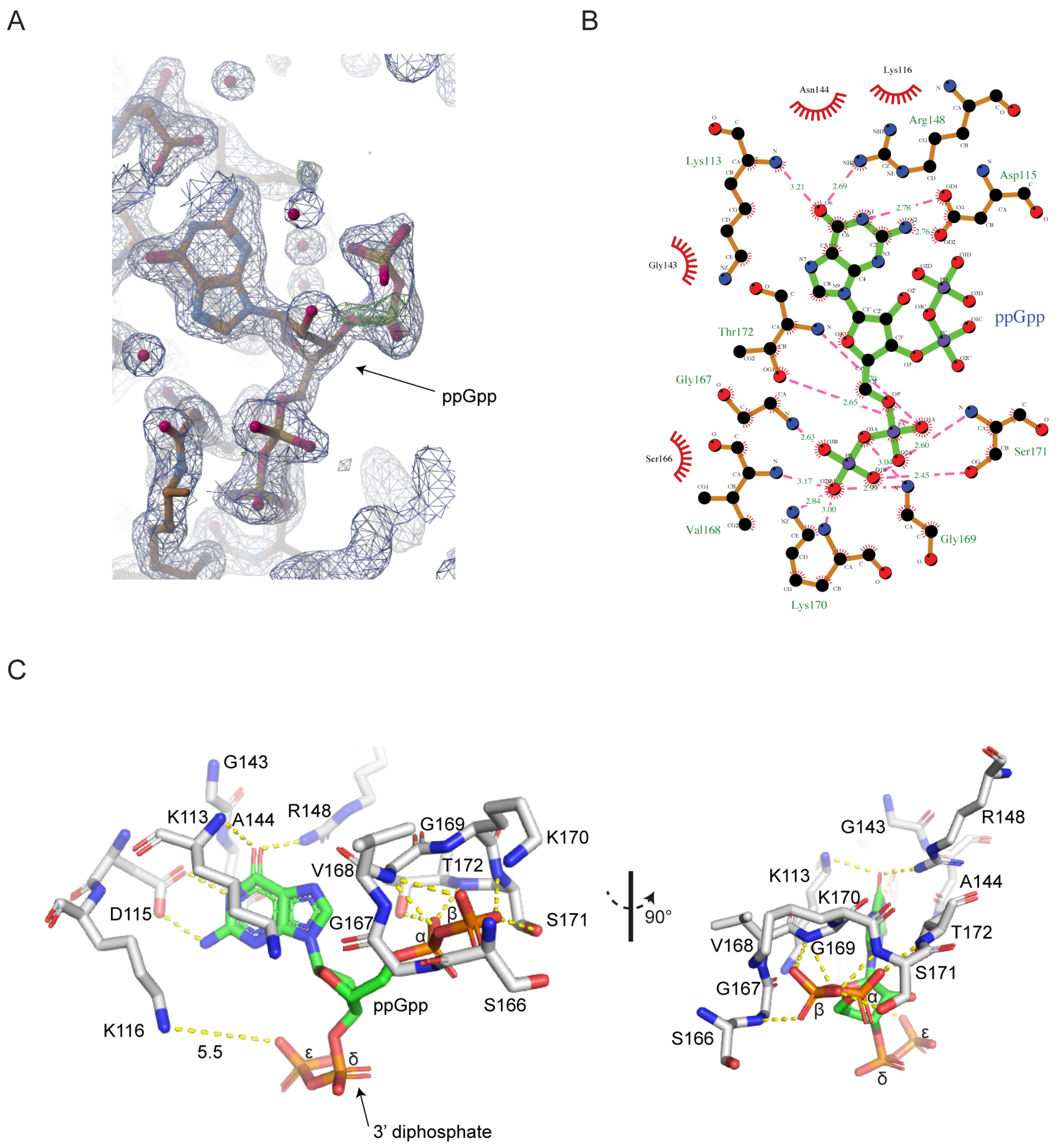


**Figure S7. The binding of ppGpp and the 30S ribosomal subunit by RsgA.** A) Fo-Fc/2Fo-Fc omit map of the ppGpp binding site of RsgA (blue mesh) overlaid with a stick model of the ppGpp ligand, coloured as follows: carbon, orange; nitrogen, blue; oxygen, red; phosphorous, yellow. Note the clear electron density due to the 3´-diphosphate of ppGpp. The Fo-Fc map is contoured at 3.1 σ and the 2Fo-Fc map is contoured at 1.6 σ. B) LigPlot maps of interacting residues involved in ppGpp binding (11). Bonds within the protein are shown in orange, whereas those within the ligand are shown in green. Hydrogen bonds are shown as pink dashed lines with their respective bond lengths indicated (Å). Protein residues are labelled and Van der Waal's contacts are represented as red curves. C) A detailed view of the ppGpp binding site of RsgA is shown as stick models. RsgA residues are represented as white models, with atoms coloured by type: carbon in green for the nucleotides and pale grey for the protein; nitrogen in blue; oxygen in red; and phosphorous in orange. Hydrogen bonds and electrostatic interactions between the protein and ligand are represented by yellow dashed lines. The uncommon bond length of the long-range stabilising interaction between K116 and the ε-phosphate of ppGpp is labelled (Å). In B) and C), Specific RsgA residues are labelled, including five of the seven G1 motif residues G167, V168, G169, K170, and S171 and two of the four G4 motif residues K113 and D115.
